# Seasonal variation in great tit (*Parus major*) energy requirements: reallocation versus increased demand

**DOI:** 10.1101/2023.04.01.535061

**Authors:** Cesare Pacioni, Marina Sentís, Catherine Hambly, John R Speakman, Anvar Kerimov, Andrey Bushuev, Luc Lens, Diederik Strubbe

## Abstract

Understanding how birds annually allocate energy to cope with changing environmental conditions and physiological states is a fundamental question in avian ecology. The two main hypotheses to explain annual patterns in energy use are “reallocation” and “increased demand”. The reallocation hypothesis suggests equal energetic costs in winter and breeding seasons, while the increased demand suggests that energy demand should be highest during breeding. Under the standard aerobic capacity model of endothermy, birds are expected to adjust the mass and/or metabolic intensity of their bodies in ways that are consistent with expected cold- and/or activity-induced costs. Here, we look for metabolic signatures of reallocation versus increased demands in the energy requirements of a small, resident passerine of a temperate-zone (great tit, *Parus major*). To do so, we measured whole-body and mass-independent basal (BMR), summit (M_sum_), and field (FMR) metabolic rates during late winter and during the chick-rearing period (breeding). We also assessed whether, and to what extent, metabolic rates conform to the predictions of the aerobic capacity model of endothermy. We found that great tits showed no substantial differences in energy expenditure between winter and the breeding season, providing support for the reallocation hypothesis. Only mass-independent M_sum_ showed seasonal variation, with significantly higher values (∼4%) in winter compared to the breeding season. Our results also lend support to the predictions of the aerobic capacity model for the evolution of endothermy, as we found that whole-body BMR and M_sum_ were positively related. We argue that both energy reallocation and the limited increase in mass-independent M_sum_ are consistent with the relatively mild winter temperatures recorded during our study period. Our results confirm that both BMR and M_sum_ are flexible traits that vary in ways that are consistent with expected cold- and/or activity-induced costs.

## Introduction

The way that birds allocate energy in order to cope with changes in environmental conditions (e.g., fluctuations in ambient temperature) and physiological states (e.g., reproductive seasonality) is a fundamental question in avian ecology. Energy expenditure is an important aspect of a species’ ecology and can vary considerably during the annual cycle. The two main hypotheses to explain annual patterns in energy use are the “reallocation hypothesis” and the “increased demand hypothesis” (Masman et al., 1988). The first hypothesis predicts a shift in energy expenditure from winter thermoregulation to reproductive activities, resulting in no net difference in seasonal energy requirements (Bryant & Tatner, 1988; Weathers & Sullivan, 1993). The second hypothesis predicts that energy demand should be highest during breeding, topping all other seasons (Gales & Green, 1990; Masman et al., 1988). Under the standard aerobic capacity model of endothermy (Bennett & Ruben, 1979), birds are expected to respond to such seasonal energetic changes in required “work” or “activity” by changing the mass and/or metabolic intensity of both their exercise and thermogenic tissues (i.e., mainly pectoral muscles) and their central organs (i.e., liver, kidney, and gut). For example, when faced with cold stress, birds can increase their pectoral muscle mass and/or metabolic intensity for improved shivering thermogenic capacity (Swanson & Vézina, 2015) and improved cold tolerance (Cooper, 2002; Schweizer et al., 2022; Swanson, 2001). Such investments in muscle mass and activity and concomitant changes in the gut and digestive organs that allow for higher daily food consumption (Chappell et al., 2011; Milbergue et al., 2018; Vézina et al., 2011) then underlie correlated changes in maximal aerobic capacity (i.e., summit metabolic rate; M_sum_) and maintenance metabolism (i.e., basal metabolic rate; BMR).

Research on tropical bird species tends to support the increased demand hypothesis. For example, Wells and Schaeffer (2012) found that M_sum_ was highest during the breeding season for seven resident bird species in Panama. The black-headed nightingale thrush (*Catharus mexicanus)* from tropical mountains upregulated its BMR in summer (∼19% increase of winter BMR; Jones et al. 2020). Other studies support the reallocation hypothesis. For example, Anava et al. (2000) documented approximately equal energy expenditure in summer and winter for non-breeding Arabian babblers (*Turdoides squamiceps*) in Israel. Magpie (*Pica pica*) (Mugaas & King, 1981), yellow-eyed junco (*Junco phaeonotus*), and dark-eyed junco (*Junco hyemalis*) (Weathers & Sullivan, 1993) similarly showed a winter energy expenditure that was not significantly different from average breeding season values. Most other studies conducted in high-to mid-latitude areas, however, tend to reject both hypotheses, as winter energy expenditure frequently exceeds that during other seasons. For example, a study on Carolina chickadees (*Poecile carolinensis*) in Ohio found that daily energy expenditure (considered as field metabolic rate, FMR) was significantly higher (∼32%) during the non-breeding season (Doherty et al., 2001), while Weathers et al. (1999) found that the energy expenditure of white-crowned sparrows (*Zonotrichia leucophrys nuttalli*) in California was 17% higher during the winter than during the breeding season. Hence, there is still debate about how generally applicable the reallocation and increased demand hypotheses are.

The aerobic capacity model is most strongly supported by a range of studies on birds living in cold areas. For instance, Dutenhoffer and Swanson (1996) found that in 10 bird species from southeast South Dakota, whole-body and mass-independent BMR and M_sum_ were positively correlated, and the same was true for black-capped chickadees (*Poecile atricapillus*) in Canada (Lewden et al., 2012). Findings from tropical birds, in contrast, found a lack of support for the aerobic capacity model, which may be explained by the fact that tropical species presumably experience less strong selection for high thermogenic capacity (Swanson & Garland, 2009; Wiersma et al., 2007a; Wiersma, 2007b; Pacioni et al., 2023). For example, Wiersma et al. (2007a) found that mass-independent M_sum_ and mass-independent BMR were not correlated in 19 tropical lowland forest birds. Moreover, M_sum_ is often considered a measure of maximal (shivering) heat production and an indicator of the level of sustainable thermogenic capacity, as many studies document a positive association with cold tolerance (Swanson, 2001; Swanson, 2010; Swanson & Liknes, 2006). However, several studies have reported that winter increases in cold tolerance can occur without corresponding changes in M_sum_ (Dawson et al., 1983; Swanson, 1993). Thus, cold tolerance and M_sum_ do not always change in tandem, and the extent of their phenotypic correlation is still controversial (Swanson et al., 2012).

Here, we look for metabolic signatures of reallocation versus increased demands in the energy requirements of a small, resident passerine of temperate zone deciduous forests (great tit, *Parus major*). We also assess whether, and to what extent, its metabolic rates conform to the predictions of the aerobic capacity model of endothermy, and how M_sum_ relates to cold tolerance. To do so, we measured the basal (BMR), summit (M_sum_), and field (FMR) metabolic rates of great tits in Belgium during late winter and during the chick-rearing period. As our study area is characterized by a maritime temperate climate with comparatively mild winters (limiting thermoregulation costs), and given the fact that great tits are single-load feeders that need to make frequent foraging trips (i.e., carrying only one prey item back to the nest), especially during the period of nestling peak food demand (Gill, 1995), we predict that (i) energy expenditure will be higher during the breeding compared to pre-breeding late winter (i.e., conforming to the increased demand hypothesis). Based on the aerobic capacity model assumptions, we further expect to find that (ii) BMR and M_sum_ are positively correlated, (iii) that body mass will be positively correlated with both BMR and M_sum_, and (iv) that M_sum_ correlates with increased cold tolerance.

## Material and methods

### Study system and fieldwork

The study was carried out in the Aelmoeseneie forest, a 39.5 ha mixed deciduous forest surrounded by residential areas and agricultural fields in Gontrode (Melle), Belgium, in February/mid-March 2022 (late winter) and in April-May 2022 (breeding). The forest has been equipped with 84 standard nest boxes for great tits since autumn 2015 (height 1.5m; dimensions 23×9×12cm; entrance 32mm) (Dekeukeleire, 2021). The climate of this region is maritime temperate, characterized by mild winters and significant precipitation in all seasons. The ambient temperature of the forest was recorded using 20 TMS-4 dataloggers, placed ∼15 cm from the ground (Wild et al., 2019). Birds were captured with both nightly nest box controls and daily mist nets. Birds were then taken to a nearby lab space, where they were ringed for individual identification, aged (1^st^ winter or adult), sexed (based on plumage characteristics), weighed to the nearest 0.1g, and kept in individual cages with food (mealworms and sunflower seeds) and water ad libitum. After the experiments, all birds were released at their original capture site. During the breeding season (starting in April), all the nest boxes were checked at least twice a week to determine occupancy by great tits and to record the day of egg laying and hatching.

### Winter BMR and breeding BMR

BMR is defined as the minimum rate of energy expenditure (measured within the thermoneutral zone of a species) that a resting, post-absorptive individual requires to maintain normothermic body temperatures (McNab, 2012). In winter, BMR was estimated by flow-through respirometry (Lighton, 2018) by measuring O_2_ consumption (VO_2_; ml/min) during the night. Prior to these measurements, 40 individuals were weighed to the nearest 0.1g and placed in airtight plastic chambers with a volume of 1.1l. All birds were food-deprived for 2 hours before respirometry. Ambient air was supplied by two pumps and divided into separate streams that were directed to a mass-flow meter (FB-8, Sable Systems) with needle valves adjusted to provide a flow of ∼400 ml/min. Then, the airstreams were directed to the eight metabolic chambers (seven for each bird and one empty reference chamber as a baseline). The excurrent airstreams were connected to a multiplexer (RM-8, Sable Systems), which allowed one chamber airstream to be sampled independently from the others. Excurrent air from the bird and the baseline channels were alternately subsampled and pulled through a Field Metabolic System (FMS-3, Sable Systems). Birds were measured alternately in cycles, together with several baselines. The time of measurement for each bird within a cycle, and the length of each cycle, depended on the number of birds within a session (around 30 minutes per bird with three cycles during the night). The average measurement time was 9 hours. All chambers were maintained inside a custom-made darkened climate control unit (Combisteel R600) set at 25°C, which is within the thermoneutral zone (TNZ) of the species. The TNZ was determined during two different nights by monitoring VO_2_ in three post-absorptive individuals during the rest phase at a series of air temperatures (15, 17.5, 20, 22.5, 25, 27.5, and 30°C), executed in a random order with ∼2 hours per each temperature (following van de Ven et al., 2013). Although the limits of the zone of thermal neutrality are not well defined by the curves (Figure 1), most individuals tended to consume less oxygen at ambient temperatures within 22.5-30°C, in line with other estimations (e.g., Bech & Mariussen, 2022). After the respirometry measurement, the birds were weighed again to the nearest 0.1g and placed back individually in their cages with water and food ad libitum. During the breeding season, 10 individuals (only females) were taken from their nest box during nighttime while roosting with their ca. 12-day-old nestlings, and taken to the lab in order to measure their BMR. The females were weighed to the nearest 0.1g and immediately transferred to the metabolic chambers. Their breeding BMR was measured as described above. At sunrise, birds were removed from the metabolic chambers, weighed to the nearest 0.1g, and returned to their original nest boxes, where they all resumed raising their offspring. We were able to measure both BMR and M_sum_ on 36 individuals.

**Figure 1.**
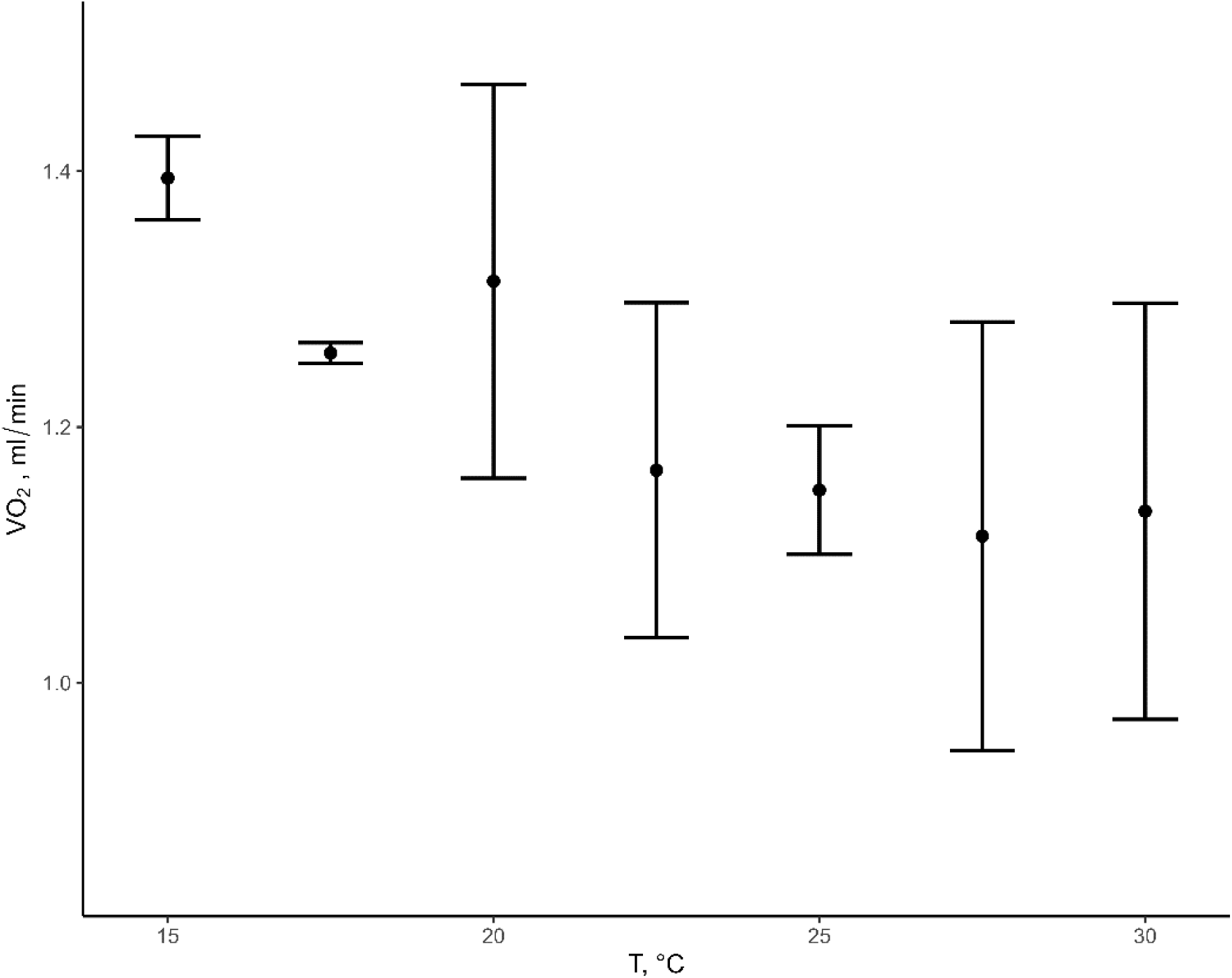
O_2_ consumption (VO_2_) expressed in ml/min at different ambient temperatures (T, °C) for great tits. The sample size for each temperature is n=3.

### Winter M_sum_ and breeding M_sum_

M_sum_ is the maximal resting metabolic rate elicited by cold exposure (Mckechnie & Swanson, 2010). In winter, M_sum_ was measured during the day after the BMR as the maximum cold-induced VO_2_ in a heliox (79% Helium-21% Oxygen) atmosphere (Rosenmann & Morrison, 1974), using the sliding cold exposure technique (Swanson et al., 1996). Prior to measurements, each bird (*n*=36) was weighed to the nearest 0.1g and placed in a 0.9l metal chamber. The chamber (with the bird) was then placed in the climate control unit, which was set at a temperature of 10°C and supplied with flowing heliox gas several minutes before starting the trial to allow the bird to acclimatize (Noakes et al., 2017). Heliox was pumped using a flow rate of ∼812 ml/min set on the FB-8. The inflow of heliox was separated into two channels: baseline and experimental. The excurrent gas stream was passed through a column of Drierite^®^ before the FMS-3 to remove water vapor. All M_sum_ trials began with a baseline heliox measurement of 7 minutes. This period allowed for the air in the metabolic chambers to be completely replaced by heliox before recording any data from the chamber. After the baseline, the setup switched to the experimental channel, and the climate control unit was set at-30°C, causing the temperature in the metabolic chamber to decrease at a rate of ∼0.8°C per minute. M_sum_ measurements were stopped whenever individuals became hypothermic, which was inferred from a steady decline in VO_2_ over several minutes (Swanson et al., 1996). After removal from the chamber, baseline values were again recorded for at least 5 minutes. We assumed a bird had reached its M_sum_ when its body temperature after a trial was ≤ 38°C (Cooper & Gessaman, 2005), measured by inserting a T-type 36 gauge thermocouple (5SC-TT-TI-36-2M; Omega) inside the cloaca. Each bird was weighed to the nearest 0.1 g both before and after metabolic tests. When the test was finished, the bird was placed back in the cage, which was located in a heated room with water and food ad libitum. During the breeding seasons, birds were captured during the day with mist nets placed at different locations in order to cover the whole forest (n=7). The individuals were then taken to the lab, weighed to the nearest 0.1g and immediately transferred to a metabolic chamber. Their M_sum_ and body temperature were measured as described above. After the measurement, the bird was again weighed and placed in a cage, which was located in a warm room with water and food ad libitum for about 30 minutes, and then immediately released at the site of capture. We decided not to measure M_sum_ on females in order to prevent them from spending too much time away from the nest. Hence, during breeding season, we do not have measurements of both BMR and M_sum_ on the same individual.

### Respirometry and data analysis

The software ExpeData (Sable Systems) was used to record trials and extract BMR (ml O_2_/min) and M_sum_ (ml O_2_/min) using equation 9.7 from Lighton (2008). The lowest stable part of the curve (average of 5 min) was selected to estimate BMR and TNZ over the entire night. M_sum_ was considered as the highest 5-min average VO_2_ over the test period, and data were corrected for drift in O_2_, CO_2_, and H_2_O baselines using the Expedata Data Drift Correction function.

### Winter FMR and breeding FMR

FMR is the total energy cost that an animal incurs during the course of a day, integrating maintenance, growth, and activity costs (e.g., associated with thermoregulation, foraging, etc.). The day after either M_sum_ or BMR measurements, FMR (kJ/day) was measured using the doubly labeled water (DLW) technique (Speakman, 1997). Blood samples were collected from 1 unlabeled individual to estimate the background isotope enrichments of ^2^H and ^18^O. Different individuals (*n*=30) were then weighed to the nearest 0.1g before being injected into the pectoral muscle with 0.1ml of DLW (661699 ppm ^18^O, 345686 ppm ^2^H). Syringes (Micro Fine + Insulin Syringe with Needle; Camlab) were weighed before and after administration (± 0.0001g) to calculate the exact dose the bird received, and the time of injection recorded. Individuals were then kept in a cloth bag for ca. 1h to allow the isotopes to mix with the body fluids. Before release at the original capture site, an initial blood sample was taken from the ulnar vein (100µl). When recaptured by nighttime nest boxes controls or daytime mist-netting 24 to 48 hours after release, another blood sample was taken from the ulnar vein in the opposite wing. The bird was weighed again to the nearest 0.1g and the time of the blood sample was recorded. All blood samples were kept in heparinized capillary tubes (Micro-Pipette 100µl; Camlab), flame-sealed using a butane torch, and stored at room temperature. Analysis of the isotopic enrichment of blood was performed blind, using a Liquid Isotope Water Analyser (Los Gatos Research, USA) (Berman et al., 2012). Initially the blood encapsulated in capillaries was vacuum distilled (Nagy, 1983), and the resulting distillate was used for analysis. Samples were run alongside five lab standards for each isotope and three International standards to correct for day to day machine variation and convert delta values to ppm. A single-pool model was used to calculate rates of CO_2_ production as recommended for use in animals less than 5 kg in body mass (Speakman, 1993). To convert CO_2_ in energy expenditure, we assumed RQ=0.75 and used 27.89 kJ/l CO_2,_ following Speakman (1997). To calculate body composition of the birds, the dilution space (Nd) was estimated from the enrichment of the deuterium in the blood sample collected 1h post injection. This reflects the total body water (TBW) content, which is then converted to fat free mass (FFM).

### Statistical analysis

Linear regression models with a Gaussian error distribution were used to test whether energy expenditure and cold tolerance (defined as the heliox temperature at which M_sum_ was reached) differed between the winter and breeding seasons, specifying FMR, BMR, and M_sum_ as dependent variables while adding sex and age as covariates. To test whether birds respond to energetic requirements primarily by adjusting the body mass and FFM of their exercise/thermoregulatory tissues and central organs, models were first run using whole-body metabolic rates. To test for changes in metabolic intensity of tissues and organs, models were then also run using mass-independent metabolic rates. Mass-independent metabolic rates are the residuals of regressions of (log) FMR, (log) BMR, and (log) M_sum_ on (log) body mass (the body mass after the trials was used). However, because no 1^st^ winter individuals were caught in the breeding season, age was only added for within-winter comparisons.

Similar linear models were used to test for positive correlations between body mass, (whole-body) metabolic rates, and cold tolerance. To avoid collinearity issues when assessing the relationship between mass-independent BMR and mass-independent M_sum_ (as mentioned above, this was only possible for the winter data as, during breeding, BMR, and M_sum_ were not measured on the same individuals), we first calculated the residuals from the regression of (log) BMR and (log) M_sum_ on (log) body mass and then used the residual values of M_sum_ as the dependent variable and the residual value of BMR as the explanatory variable (Downs et al., 2013). As body fat is metabolically inactive, we also used FFM obtained through deuterium dilution instead of body mass. We then conducted a similar analysis on a subset for whom FFM data was available (n=16; 10 in late winter and 6 in breeding season).

For all models, we used a backward stepwise procedure to eliminate non-significant interactions and variables. Post-hoc comparisons between species and seasons were performed with the emmeans function in the ‘emmeans’ package (Lenth, 2019). We used interquartile ranges as a criterion to identify outliers by using the quantile function. Then, we used the subset() function to eliminate outliers. For all models, the normality of residuals was tested and verified (i.e., Shapiro-Wilk W > 0.9), and the significance level was set at p ≤ 0.05. Body mass, FMR, BMR, and M_sum_ were log transformed before all analyses. Statistical analysis was performed using R software v. 4.2.2 (R Core Team 2022). Details of the statistical analysis are available in Supplementary file (RMarkdown HTML).

## Results

During winter, we measured BMR of 40 individuals (19 males and 21 females, including 35 adults and 5 1^st^ winter). For all but four of these individuals, we also measured their winter M_sum_ (16 males and 20 females, including 31 adults and 5 1^st^ winter). During the breeding season, we measured BMR of ten adult females and M_sum_ of six adult males and one adult female. Due to difficulties in recapturing individuals within 24 to 48 hours for the measurement of FMR, we only obtained five measurements (two during winter and three during breeding; 1 male adult and 4 female adults).

Great tits showed no difference in proxies of energy expenditure between winter and breeding, as whole-body FMR, BMR, and M_sum_ did not differ between the two seasons (all p>0.1). Regarding mass-independent metabolic rates, only M_sum_ was ∼4% higher in winter than during the breeding season (p<0.01). Great tits were able to significantly tolerate colder temperatures during the winter compared to the breeding season (all p<0.05), with a winter average of-18.1 ± 1.7 °C and a breeding average of-16.5 ± 1.5 °C (heliox temperatures at which M_sum_ was reached). Male great tits (both in late winter and during the breeding season) were significantly heavier (17.2 ± 1.4 g) than females (15.7 ± 1.0 g) (p<0.001), which was associated with a higher whole-body M_sum_ (p<0.05). The body mass after BMR measurements was considered. 1^st^ winter individuals had a significantly higher body mass than adults (p<0.05), which was associated with a higher whole-body and mass-independent M_sum_ (p<0.05). The average metabolic expansibility (ME, the ratio of M_sum_ over BMR) during the winter season was 3.94, while during the breeding season the average was 3.19 (population level, see above, as we did not gather BMR and M_sum_ on the same individuals during the breeding season). BMR (both whole-body and mass-independent) was not significantly correlated with the number of chicks nor with the laying date. Table 1 summarizes sample sizes, means, standard deviations (SD), minimum (Min), and maximum (Max) of body mass, FMR, BMR, M_sum_, and cold tolerance for both seasons.

**Table 1.** Mean ± standard deviation (SD), minimum (Min) - maximum (Max) values for FMR_w_ (late winter), FMR_b_ (breeding), BMR_w_ (late winter), BMR_b_ (breeding), M_sum__w_ (late winter), M_sum__b_ (breeding), cold tolerance (late winter and breeding) and body mass (after BMR measurements; late winter and breeding) per sex and age. Values with * refer to only one individual.

|  |  | Male |  | Female |  | Adult |  | 1st winter |  |
| --- | --- | --- | --- | --- | --- | --- | --- | --- | --- |
| | | Mean ± SD | Min - Max | Mean ± SD | Min - Max | Mean ± SD | Min - Max | Mean ± SD | Min - Max |
| Whole-body | FMR<sub>w</sub> [kJ/day] | 67.4* | | 65.2* | | 66.3 ± 1.5 | 65.2 - 67.4 | - | - |
| | FMR<sub>b</sub> [kJ/day] | - | - | 72.7 ± 21.1 | 56.3 - 96.5 | 72.7 ± 21.1 | 56.3 - 96.5 | - | - |
| | BMR<sub>w</sub> [ml O<sub>2</sub>/min] | 1.23 ± 0.23 | 0.86 - 1.89 | 1.20 ± 0.18 | 0.81 - 1.60 | 1.22 ± 0.20 | 0.81 - 1.89 | 1.32 ± 0.14 | 1.22 - 1.56 |
| | BMR<sub>b</sub> [ml O<sub>2</sub>/min] | - | - | 1.35 ± 0.25 | 0.89 - 1.58 | 1.35 ± 0.25 | 0.89 - 1.58 | - | - |
| | Msum<sub>w</sub> [ml O<sub>2</sub>/min] | 4.94 ± 0.58 | 3.92 - 6.75 | 4.27 ± 0.58 | 2.61 - 5.33 | 4.57 ± 0.66 | 2.61 - 6.75 | 5.22 ± 0.33 | 4.83 - 5.70 |
| | Msum<sub>b</sub> [ml O<sub>2</sub>/min] | 4.38 ± 0.52 | 3.92 - 5.21 | 3.84* | | 4.29 ± 0.52 | 3.84 - 5.21 | - | - |
| Mass-independent<br>[residuals] | FMR<sub>w</sub> | 0.06* | | -0.11* | | -0.02 ± 0.12 | -0.11 - 0.06 | - | - |
| | FMR<sub>b</sub> | - | - | 0.02 ± 0.17 | -0.15 - 0.20 | 0.02 ± 0.17 | -0.15 - 0.20 | - | - |
| | BMR<sub>w</sub> | -0.02 ± 0.13 | -0.29 - 0.33 | -0.01 ± 0.10 | -0.21 - 0.20 | -0.02 ± 0.17 | -0.15 - 0.19 | 0.06 ± 0.23 | -0.16 - 0.33 |
| | BMR<sub>b</sub> | - | - | 0.07 ± 0.17 | -0.21 - 0.24 | 0.07 ± 0.17 | -0.21 - 0.24 | - | - |
| | Msum<sub>w</sub> | 0.05 ± 0.12 | -0.07 - 0.38 | -0.01 ± 0.13 | -0.42 - 0.18 | 0.04 ± 0.14 | -0.07 - 0.38 | 0.08 ± 0.07 | -0.00 - 0.15 |
| | Msum<sub>b</sub> | -0.09 ± 0.11 | -0.23 - 0.07 | -0.16* | | -0.01 ± 0.11 | -0.23 - 0.07 | - | - |
| Cold tolerance<sub>w</sub> [°C] | | -18.5 ± 1.6 | -22.0 - -15.0 | -17.9 ± 1.8 | -20.0 - -14.0 | -18.1 ± 1.7 | -22.0 - -14.0 | -18.0 ± 1.1 | -19.2 - -17.0 |
| Cold tolerance<sub>b</sub> [°C] | | -16.6 ± 1.7 | -19.0 - -15.0 | -16.0* | | -16.5 ± 1.5 | -19.0 - -15.0 | - | - |
| Body mass<sub>w</sub> [g] | | 18.0 ± 1.4 | 14.9 - 20.4 | 16.3 ± 1.1 | 14.5 - 18.3 | 17.1 ± 1.5 | 14.5 - 20.4 | 18.4 ± 1.2 | 17.0 - 20.1 |
| Body mass<sub>b</sub> [g] | | 17.6 ± 0.9 | 16.6 - 18.8 | 16.8 ± 0.9 | 15.8 - 18.4 | 17.1 ± 0.9 | 15.8 - 18.8 | - | - |

We found evidence for a positive correlation between (log) body mass and both (log) BMR (p<0.01) and (log) M_sum_ (p<0.05) (Figure 2). Moreover, (log) BMR and (log) M_sum_ were also positively correlated (p<0.05), and (log) M_sum_ was higher for individuals who could stand lower temperatures (i.e., had higher cold tolerance) (p<0.01).

**Figure 2.**
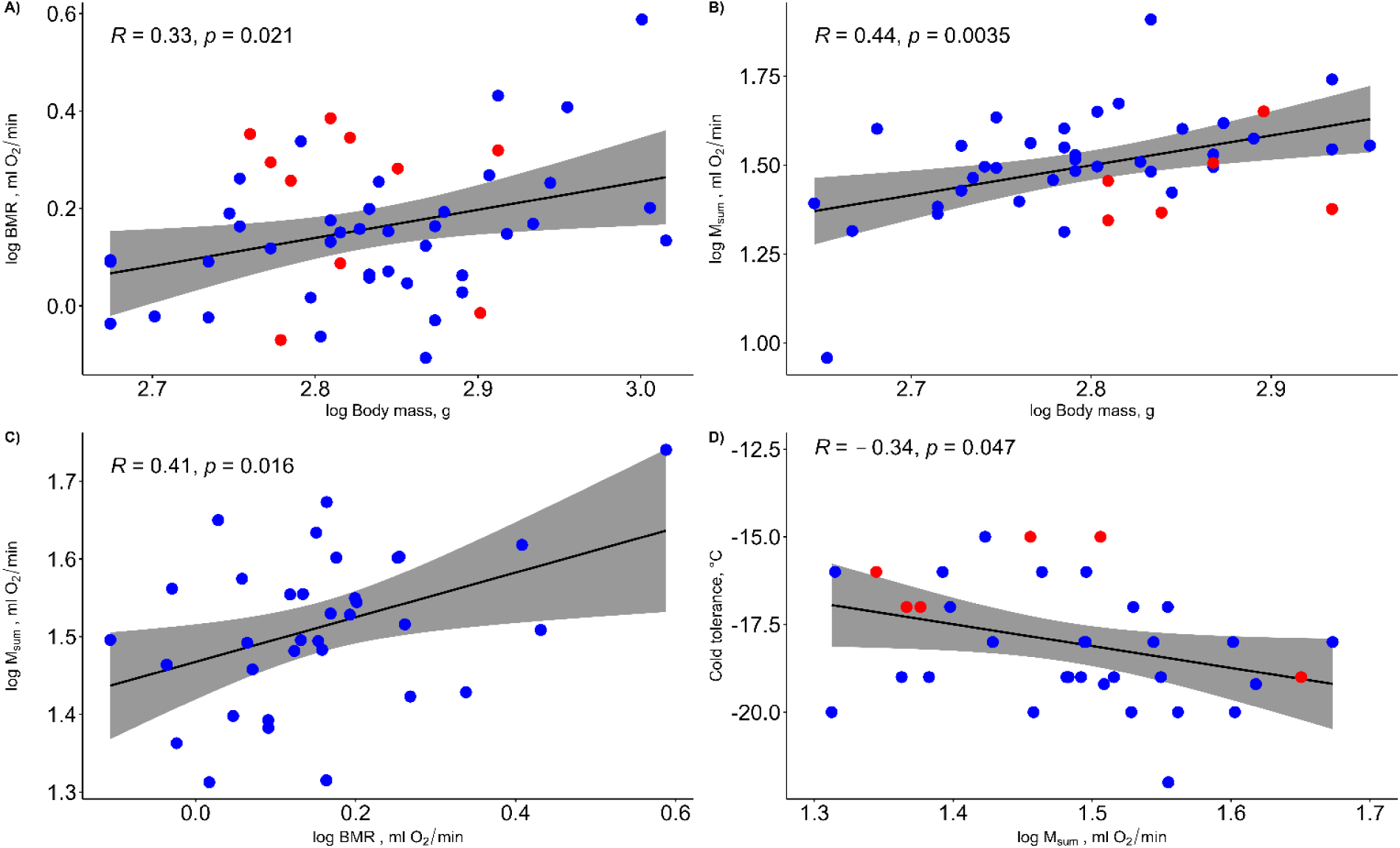
A) The relationship between (log) BMR and (log) body mass. B) The relationship between (log)M_sum_ and (log) body mass. C) The relationship between log(M_sum_) and log(BMR). D) The relationship between cold tolerance (heliox temperature at which M_sum_ was reached) and log(M_sum_). Blue dots = winter. Red dots = breeding season.

When we substituted (log) body mass with (log) FFM, the results showed no correlation between (log) FFM, (log) BMR or (log) M_sum_. We also investigated the mass-independent metabolic rates by regressing (log) FMR, (log) BMR, and (log) M_sum_ on (log) FFM to calculate the residuals. These mass-independent metabolic rates did not exhibit significant correlation and did not vary with seasonal changes. However, the sample size was relatively small, with only 16 individuals having FFM data (10 in late winter and 6 in the breeding season), which may have restricted the detection of variations and correlations. The analysis showed that the only significant difference observed when using FFM was that individuals with higher cold tolerance exhibited higher mass-independent M_sum_ (i.e., those who could tolerate lower temperatures; p<0.01). Nonetheless, when we used body mass instead of FFM in the same subset of individuals (n=16), we still observed a significant relationship between mass-independent M_sum_ and cold tolerance (p<0.001).

## Discussion

The aim of this study was to test the ‘reallocation’ and ‘increased demand’ hypotheses on great tits living in a temperate area and to assess whether predictions from the aerobic model of endothermy underlie (changes in) energy expenditure and cold tolerance between seasons. The increased demand hypothesis predicts that energy demand should be highest during breeding, while the reallocation hypothesis expects no net difference in seasonal energy requirements because of a shift in energy expenditure from winter thermoregulation to reproductive activities in spring. Overall, our analysis of the whole-body metabolism of great tits revealed that there were no substantial differences in energy expenditure between the winter and the breeding season, providing support for the reallocation hypothesis. Only mass-independent M_sum_ showed seasonal variation, with significantly higher values in winter compared to the breeding season, likely reflecting an adjustment to tolerate colder winter temperatures. Cold tolerance in winter was indeed positively related to M_sum_. Our results lend support to the predictions of the aerobic capacity model for the evolution of endothermy, as we found that whole-body BMR and M_sum_ are positively related and that both are positively related to body mass.

### Reallocation versus increased demand hypothesis

Studies focusing on small passerines at higher latitudes or in continental areas with harsher winter climates often reject both the increased demand and reallocation hypotheses because of a higher winter energy expenditure (Cooper, 2000; Doherty et al., 2001; Weathers et al., 1999; Webster & Weathers, 2000). Increased metabolic rates and body masses allow small passerines to withstand cold exposure for longer time periods (Piersma, 1984; Stuebe & Ketterson, 1982) and are thus a key strategy for increasing winter survival in cold areas (Broggi et al., 2003). Given the comparatively mild winters in Belgium, we had expected that the breeding season would be more demanding for great tits, yet our data show no substantial increase in metabolic rates or body mass from late winter to the breeding season, and the results thus lend support to the reallocation hypothesis. It is indeed surprising that late winter is as energetically challenging as the chick-rearing period, especially as ambient temperatures during February and March 2022, when our pre-breeding period measurements were carried out, were even higher than typical. Temperature averages from the forest loggers were 6.5 ± 1.8 °C in February and 9.3 ± 2.9 °C in March, compared to a normal February average temperature of about 4.2 °C and about 7.1 °C for March (Royal Meteorological Institute; RMI). The fact that great tits are relatively small birds (∼16.4g, Table 1) may offer a potential explanation here, as they can rapidly lose body heat because of their high surface-to-volume ratios (Cooper, 2000; Dawson et al., 1983; Doherty et al., 2001; Weathers et al., 1999; Webster & Weathers, 2000), potentially resulting in significant thermoregulatory demands even in milder climates.

The finding that energy expenditure in our study population corroborates the reallocation hypothesis contrasts with the results of several great tit studies assessing seasonal variation in BMR and body mass. For example, a study of geographical and seasonal variation in BMR in great tits showed that BMR peaked during midwinter (Broggi et al., 2019) for two populations in Oulu (Finland) and Lund (Sweden). A study focusing on body mass regulation in great tits in north-eastern Europe found an increase in body mass under conditions of extremely low ambient temperatures (i.e., minus 37°C, Krams et al., 2010). Similar findings have been reported for other small-bodied species, such as the Eurasian tree sparrow (*Passer montanus*) in China, for which Zheng et al. (2008) showed a higher BMR in winter when ambient temperatures averaged ∼-15 °C. The contrasting findings in our study population are thus likely due to significantly warmer winter temperatures compared to many other studies of seasonal variation in energy expenditure in small passerines.

To our knowledge, our study is the first to measure seasonal variation of M_sum_ in great tits, and while all other proxies of energy expenditure studied here were generally similar in late winter versus the breeding season, we did find evidence for a small but statistically significant (∼4%) increase in mass-independent M_sum_ during winter. A winter increase in mass-independent M_sum_ is a common finding in many studies of temperate-zone species (Mckechnie & Swanson, 2010), though it is typically higher than the increase found here (i.e., 10 to 50% higher; Swanson, 2010). Birds can achieve higher metabolic heat production and thus higher cold tolerance either by increasing their shivering thermogenic capacity by investing in large (mainly pectoral) muscle mass (see below) or, alternatively, by elevating the metabolic intensity of their tissues (Petit et al., 2014, 2017). Mass-independent M_sum_ changes represent changes in the heat production per unit tissue mass (McKechnie, 2008; Swanson, 2010), and have been linked to adjustments in tissue mitochondrial density, upregulation of avian mitochondrial uncoupling proteins (Dridi et al., 2004) or increased activity of oxidative enzymes (Swanson 2010). Such changes may allow birds to rapidly respond to changes in weather conditions, such as the arrival of cold spells, without requiring the synthesis of new shivering tissues (Vézina et al., 2011). The rather limited increase in M_sum_ documented here is consistent with the relatively mild winter temperatures recorded during our study period, but more research is needed to test the relative importance of shivering versus non-shivering thermogenesis in birds exposed to different climates and weather regimens.

Compared to other studies on the energetic metabolism of great tits in Europe, we found that our study birds were characterized by substantially lower metabolic rates and body masses (Supplementary material; Table S1 and S2). However, mass-specific BMR and FMR values in this study fall within the range of reported values (Table S1 and S2). The widespread presence of bird feeders in and near our study area (Dekeukeleire, 2021) implies that birds probably had easy access to constant and reliable food sources. This may have enabled them to trade off starvation and predation risks by reducing their body mass in order to increase their chances of escaping predators (Lima, 1986). More fine-grained comparisons between our study and others are hampered because of our smaller sample sizes, a number of methodological differences (e.g., the use of open versus closed respirometry for BMR, intramuscular injection of doubly-labeled water in the pectoralis major versus intraperitoneal injection), and differences in measured life-history covariates (such as clutch size). More research, including meta-analyses summarizing longitudinal studies on the winter and the breeding energetics of individuals and populations, is needed to fully unravel the factors that influence energy allocation in wild birds and how these may vary across latitude and between habitats.

### Aerobic capacity model

We found that birds characterized by a higher BMR also had higher M_sum_ values, but that the metabolic intensity of central organs was not correlated to the metabolic intensity of thermogenic and exercise tissues. These results lend at least partial support to the aerobic capacity model of endothermy, which postulates a positive correlation between minimum (BMR) and maximum (M_sum_) metabolic output. The fact that only whole-body mass metabolic rates conformed to expectations suggests that correlations were mainly driven by variation in body mass. This suggests that great tits manage energy demands primarily by adjusting the size of their central organs (which determine BMR) and the mass of their exercise and thermoregulatory tissues (which determine M_sum_), rather than by their metabolic intensity. Interspecific comparisons of BMR and M_sum_ often found correlations between both whole-body and mass-independent basal and maximal rates, at least for non-tropical birds (Dutenhoffer & Swanson, 1996; Rezende et al., 2002), though intraspecific studies have found results similar to ours. For example, Swanson et al. (2012) showed that whole-body BMR and M_sum_ were significantly positively correlated for black-capped chickadees and house sparrows (*Passer domesticus*) from South Dakota (USA), but mass-independent rates were not. Vézina et al. (2006) found the same for overwintering red knots (*Calidris canutus*) in the Netherlands. The absence of a mass-independent correlation may be due to the fact that body mass differences between individuals can reflect differences in body composition, such as in (metabolically relatively inert) fat mass, while scaling metabolic rates by body mass to obtain mass-independent rates assumes a constant contribution of mass to metabolic rates (Daan et al., 1990; Hayes & Shonkwiler, 1996). Because body composition adjustment is a prominent mechanism underlying metabolic flexibility in birds (Swanson, 2010), whole-body metabolic rates are generally believed to be most informative for intraspecific analyses of avian energetics (Swanson et al., 2012).

As expected, birds with a higher M_sum_ were able to tolerate colder temperatures both in the pre-breeding late winter period and during chick rearing, suggesting that pectoral muscle-driven shivering thermogenesis constitutes a main mechanism of homeothermy under cold stress in small birds. In birds, the capacity for sustaining flight (i.e., parental care activity) and for shivering thermogenesis (i.e., temperature regulation) are both functions of skeletal muscle mass (Guglielmo, 2010). During the late winter period, high M_sum_ enabled great tits to tolerate colder temperatures, which conforms to several inter- and intraspecific studies on avian cold tolerance. For example, Cooper (2000) found that higher winter pectoral muscle masses resulted in higher M_sum_ and an increased ability to sustain cold temperatures in the mountain chickadee (*Poecile gambeli*) and the juniper titmouse (*Baeolophus ridgwayi*) in North America. Likewise, Liknes and Swanson (2011) showed that the pectoral muscle mass acts as a significant contributor to M_sum_ values, in turn increasing thermogenic capacity and cold tolerance for several South Dakota (USA) passerines. During the breeding period, there is little need for shivering thermogenesis, but sustained muscle activity is required for powering parental care activities, such as foraging flights (Koteja, 2000). Above, we found that, in line with the reallocation hypothesis, whole-body M_sum_ did not substantially differ between late winter and the breeding period. Thus, our study birds likely had broadly similar muscle mass in both periods, explaining the similar degree of cold tolerance, with the slightly higher winter cold tolerance (1.5°C colder) likely due to slightly higher winter M_sum_ intensity (see above). Thus, our results corroborate the literature supporting the view that M_sum_ and BMR are flexible traits that vary in ways that are consistent with expected cold- and/or activity-induced costs (such as thermoregulation and parental care).

## Conclusions

In conclusion, our findings lend support to the reallocation hypothesis and the aerobic model of endothermy. We only found a very modest (4%), but statistically significant, increase in mass-independent M_sum_, and mass-independent M_sum_ was positively correlated with cold tolerance. We argue that both energy reallocation and the limited increase in mass-independent M_sum_ are consistent with the relatively mild winter temperatures recorded during our study period. Our results suggest that both BMR and M_sum_ are flexible traits that vary in ways that are consistent with expected cold- and/or activity-induced costs.

## Funding

This study was supported by the Research Foundation – Flanders (grant #G0E4320 N), by Methusalem Project 01M00221 (Ghent University), under bilateral research cooperation with the Russian Science Foundation (grant #20-44-01005).

## Author contributions

Cesare Pacioni: Conceptualization, Methodology, Validation, Formal analysis, Investigation, Data curation, Writing – original draft, Visualization. Marina Sentís: Conceptualization, Methodology, Investigation, Writing – review & editing. Catherine Hambly: Methodology, Validation, Formal analysis, Writing – review & editing. John R Speakman: Methodology, Validation, Formal analysis, Writing – review & editing. Anvar Kerimov: Conceptualization, Methodology, Validation, Writing – review & editing. Andrey Bushuev: Conceptualization, Methodology, Validation, Writing – review & editing. Luc Lens: Conceptualization, Validation, Writing – review & editing, Supervision. Diederik Strubbe: Conceptualization, Methodology, Validation, Investigation, Writing – review & editing, Visualization, Supervision.

## Conflict of Interest

There were no conflicts of interest.

## Data availability

The data used in this manuscript can be found at (https://data.mendeley.com/datasets/fk83s6j4x2) while the statistical scripts used can be consulted via Supplementary file 1 (RMarkdown HTML).

## Acknowledgements

We thank ForNaLab for providing us with access to their facilities. Especially we would like to thank Luc Willems for his support and Dries Landuyt for providing us with temperature data from the dataloggers. In addition, we are thankful for the assistance provided by bachelor student Hanne Danneels during the data collection process. The authors would like to express sincere gratitude to An Martel for her valuable instruction and guidance in teaching the techniques of injection and blood collection. This study acknowledges funding by FWO-Vlaanderen (project G0E4320N), by Methusalem Project 01M00221 (Ghent University) awarded to Frederick Verbruggen, Luc Lens and An Martel, and by the Russian Science Foundation (project 20-44-01005). Marina Sentís acknowledges the support of FWO-Vlaanderen (project 11E1623N).

## Notes

https://data.mendeley.com/datasets/fk83s6j4x2

### Competing Interest Statement

The authors have declared no competing interest.

