## Supplementary file 1 for "Seasonal variation in great tit (*Parus major*) energy requirements: reallocation versus increased demand"

###### Pacioni et al. 2023

### 1 Statistical analysis

### 2 Aerobic capacity model

### 3 BMR ~ Msum

#### 3.1 Mass-independent

```
ggscatter(Data_NoOL, x = "resBMR", y = "resMsum", 
          add = "reg.line", conf.int = TRUE, 
          cor.coef = TRUE, cor.method = "pearson",
          xlab = "mass-independent BMR", ylab = "mass-independent Msum")
```

```
#FFM as body mass

ggscatter(Data_FFM, x = "resBMR_FFM", y = "resMsum_FFM", 
          add = "reg.line", conf.int = TRUE, 
          cor.coef = TRUE, cor.method = "pearson",
          xlab = "mass-independent BMR", ylab = "mass-independent Msum")
```

#### 3.2 Whole-body

```
ggscatter(Data_NoOL, x = "logBMR", y = "logMsum", 
          add = "reg.line", conf.int = TRUE, 
          cor.coef = TRUE, cor.method = "pearson",
          xlab = "(log)BMR [ml/min]", ylab = "(log)Msum [ml/min]")
```

### 4 BMR - Msum ~ Body mass

```
ggscatter(Data_NoOL, x = "logmassBMR_1", y = "logBMR", 
          add = "reg.line", conf.int = TRUE, 
          cor.coef = TRUE, cor.method = "pearson",
          xlab = "(log)Body Mass [g]", ylab = "(log)BMR [ml/min]")
```

```
a<-lm(logBMR~logmassBMR_1,Data_FFM)
summary(a)
```

```
## 
## Call:
## lm(formula = logBMR ~ logmassBMR_1, data = Data_FFM)
## 
## Residuals:
##       Min        1Q    Median        3Q       Max 
## -0.263620 -0.086446  0.000007  0.097149  0.238960 
## 
## Coefficients:
##              Estimate Std. Error t value Pr(>|t|)
## (Intercept)   0.22683    1.99488   0.114    0.911
## logmassBMR_1 -0.01183    0.70136  -0.017    0.987
## 
## Residual standard error: 0.1506 on 13 degrees of freedom
##   (1 observation deleted due to missingness)
## Multiple R-squared:  2.19e-05,   Adjusted R-squared:  -0.0769 
## F-statistic: 0.0002847 on 1 and 13 DF,  p-value: 0.9868
```

```
AIC(a)
```

```
## [1] -10.36356
```

```
a_1<-lm(logBMR~logFFM,Data_FFM)
summary(a_1)
```

```
## 
## Call:
## lm(formula = logBMR ~ logFFM, data = Data_FFM)
## 
## Residuals:
##      Min       1Q   Median       3Q      Max 
## -0.27169 -0.09020  0.00466  0.10250  0.23726 
## 
## Coefficients:
##             Estimate Std. Error t value Pr(>|t|)
## (Intercept)   0.6155     1.5437   0.399    0.697
## logFFM       -0.1533     0.5601  -0.274    0.789
## 
## Residual standard error: 0.1502 on 13 degrees of freedom
##   (1 observation deleted due to missingness)
## Multiple R-squared:  0.005729,   Adjusted R-squared:  -0.07075 
## F-statistic: 0.07491 on 1 and 13 DF,  p-value: 0.7886
```

```
AIC(a_1)
```

```
## [1] -10.44941
```

```
#FFM as body mass
ggscatter(Data_FFM, x = "logFFM", y = "logBMR", 
          add = "reg.line", conf.int = TRUE, 
          cor.coef = TRUE, cor.method = "pearson",
          xlab = "FFM [g]", ylab = "(log)BMR [ml/min]")
```

```
ggscatter(Data_NoOL, x = "logmassMsum_1", y = "logMsum", 
          add = "reg.line", conf.int = TRUE, 
          cor.coef = TRUE, cor.method = "pearson",
          xlab = "(log)BodyMass [g]", ylab = "(log)Msum [ml/min]")
```

```
#FFM as body mass
ggscatter(Data_FFM, x = "logFFM", y = "logMsum", 
          add = "reg.line", conf.int = TRUE, 
          cor.coef = TRUE, cor.method = "pearson",
          xlab = "FFM [g]", ylab = "(log)Msum [ml/min]")
```

```
a<-lm(logMsum~logmassMsum_1,Data_FFM)
summary(a)
```

```
## 
## Call:
## lm(formula = logMsum ~ logmassMsum_1, data = Data_FFM)
## 
## Residuals:
##      Min       1Q   Median       3Q      Max 
## -0.20468 -0.05864 -0.02836  0.04252  0.35289 
## 
## Coefficients:
##               Estimate Std. Error t value Pr(>|t|)
## (Intercept)     0.7532     3.2965   0.228    0.824
## logmassMsum_1   0.2834     1.1754   0.241    0.815
## 
## Residual standard error: 0.1526 on 9 degrees of freedom
##   (5 observations deleted due to missingness)
## Multiple R-squared:  0.006416,   Adjusted R-squared:  -0.104 
## F-statistic: 0.05812 on 1 and 9 DF,  p-value: 0.8149
```

```
AIC(a)
```

```
## [1] -6.349178
```

```
a_1<-lm(logMsum~logFFM,Data_FFM)
summary(a_1)
```

```
## 
## Call:
## lm(formula = logMsum ~ logFFM, data = Data_FFM)
## 
## Residuals:
##      Min       1Q   Median       3Q      Max 
## -0.19277 -0.04873 -0.02833  0.03610  0.35139 
## 
## Coefficients:
##             Estimate Std. Error t value Pr(>|t|)
## (Intercept)   0.8740     2.7733   0.315    0.760
## logFFM        0.2448     1.0073   0.243    0.813
## 
## Residual standard error: 0.1526 on 9 degrees of freedom
##   (5 observations deleted due to missingness)
## Multiple R-squared:  0.006519,   Adjusted R-squared:  -0.1039 
## F-statistic: 0.05905 on 1 and 9 DF,  p-value: 0.8134
```

```
AIC(a_1)
```

```
## [1] -6.350315
```

### 5 BMR ~ Body mass (Winter)

```
ggscatter(BMR_w, x = "logmassBMR_1", y = "logBMR", 
          add = "reg.line", conf.int = TRUE, 
          cor.coef = TRUE, cor.method = "pearson",
          xlab = "(log)BodyMass [g]", ylab = "(log)BMR [ml/min]")
```

```
# FFM
ggscatter(Data_FFM_W, x = "logFFM", y = "logBMR", 
          add = "reg.line", conf.int = TRUE, 
          cor.coef = TRUE, cor.method = "pearson",
          xlab = "(log)FFM [g]", ylab = "(log)BMR [ml/min]")
```

### 6 BMR ~ Body mass (Breeding)

```
ggscatter(BMR_b, x = "logmassBMR_1", y = "logBMR", 
          add = "reg.line", conf.int = TRUE, 
          cor.coef = TRUE, cor.method = "pearson",
          xlab = "(log)BodyMass [g]", ylab = "(log)BMR [ml/min]")
```

### 7 Msum ~ Body mass (Winter)

```
ggscatter(BMR_w, x = "logmassMsum_1", y = "logMsum", 
          add = "reg.line", conf.int = TRUE, 
          cor.coef = TRUE, cor.method = "pearson",
          xlab = "(log)BodyMass [g]", ylab = "(log)BMR [ml/min]")
```

#Body mass ~ BMR (Breeding)

```
ggscatter(BMR_b, x = "logmassMsum_1", y = "logMsum", 
          add = "reg.line", conf.int = TRUE, 
          cor.coef = TRUE, cor.method = "pearson",
          xlab = "(log)BodyMass [g]", ylab = "(log)BMR [ml/min]")
```

#Seasonal variations: reallocation vs increase demand

#Body mass ~ seasons + age

```
BodyMass_mod<-lm(logmassBMR_1 ~ season+age+sex+age:sex, data=Data_BMR)
summary(BodyMass_mod)
```

```
## 
## Call:
## lm(formula = logmassBMR_1 ~ season + age + sex + age:sex, data = Data_BMR)
## 
## Residuals:
##       Min        1Q    Median        3Q       Max 
## -0.177781 -0.041515 -0.002651  0.044079  0.136393 
## 
## Coefficients:
##             Estimate Std. Error t value Pr(>|t|)    
## (Intercept)  2.82071    0.02234 126.265  < 2e-16 ***
## seasonw     -0.03305    0.02736  -1.208 0.233373    
## agejuv       0.08591    0.07239   1.187 0.241566    
## sexM         0.09148    0.02413   3.791 0.000444 ***
## agejuv:sexM -0.04524    0.08259  -0.548 0.586516    
## ---
## Signif. codes:  0 '***' 0.001 '**' 0.01 '*' 0.05 '.' 0.1 ' ' 1
## 
## Residual standard error: 0.07064 on 45 degrees of freedom
## Multiple R-squared:  0.3213, Adjusted R-squared:  0.2609 
## F-statistic: 5.325 on 4 and 45 DF,  p-value: 0.001345
```

```
BodyMass_mod<-lm(logmassBMR_1 ~ season+age+sex, data=Data_BMR)
summary(BodyMass_mod)
```

```
## 
## Call:
## lm(formula = logmassBMR_1 ~ season + age + sex, data = Data_BMR)
## 
## Residuals:
##       Min        1Q    Median        3Q       Max 
## -0.175574 -0.042006 -0.002316  0.043133  0.138600 
## 
## Coefficients:
##             Estimate Std. Error t value Pr(>|t|)    
## (Intercept)  2.82071    0.02217 127.236  < 2e-16 ***
## seasonw     -0.03140    0.02699  -1.163 0.250663    
## agejuv       0.05115    0.03458   1.479 0.145935    
## sexM         0.08762    0.02290   3.826 0.000391 ***
## ---
## Signif. codes:  0 '***' 0.001 '**' 0.01 '*' 0.05 '.' 0.1 ' ' 1
## 
## Residual standard error: 0.0701 on 46 degrees of freedom
## Multiple R-squared:  0.3167, Adjusted R-squared:  0.2722 
## F-statistic: 7.108 on 3 and 46 DF,  p-value: 0.0005065
```

```
BodyMass_mod<-lm(logmassBMR_1 ~ season+sex, data=Data_BMR)
summary(BodyMass_mod)
```

```
## 
## Call:
## lm(formula = logmassBMR_1 ~ season + sex, data = Data_BMR)
## 
## Residuals:
##       Min        1Q    Median        3Q       Max 
## -0.186341 -0.047211  0.000042  0.051408  0.127833 
## 
## Coefficients:
##             Estimate Std. Error t value Pr(>|t|)    
## (Intercept)  2.82071    0.02245 125.658  < 2e-16 ***
## seasonw     -0.02896    0.02727  -1.062    0.294    
## sexM         0.09595    0.02248   4.269 9.45e-05 ***
## ---
## Signif. codes:  0 '***' 0.001 '**' 0.01 '*' 0.05 '.' 0.1 ' ' 1
## 
## Residual standard error: 0.07099 on 47 degrees of freedom
## Multiple R-squared:  0.2842, Adjusted R-squared:  0.2538 
## F-statistic: 9.332 on 2 and 47 DF,  p-value: 0.0003864
```

```
shapiro.test(resid(BodyMass_mod))
```

```
## 
##  Shapiro-Wilk normality test
## 
## data:  resid(BodyMass_mod)
## W = 0.98006, p-value = 0.5544
```

```
plot(allEffects(BodyMass_mod))
```

```
plot_model(BodyMass_mod, type="emm", terms=c("season"), mdrt.values="meansd", ppd=TRUE, line.size=1,dot.size=5)
```

```
emmeans(BodyMass_mod, pairwise ~ season, nesting=NULL)
```

```
## $emmeans
##  season emmean     SE df lower.CL upper.CL
##  b        2.87 0.0251 47     2.82     2.92
##  w        2.84 0.0112 47     2.82     2.86
## 
## Results are averaged over the levels of: sex 
## Confidence level used: 0.95 
## 
## $contrasts
##  contrast estimate     SE df t.ratio p.value
##  b - w       0.029 0.0273 47   1.062  0.2937
## 
## Results are averaged over the levels of: sex
```

```
BodyMass_age_mod<-lm(logmassBMR_1 ~ age, data=Data_BMR)
summary(BodyMass_age_mod)
```

```
## 
## Call:
## lm(formula = logmassBMR_1 ~ age, data = Data_BMR)
## 
## Residuals:
##       Min        1Q    Median        3Q       Max 
## -0.151350 -0.051352 -0.007104  0.047138  0.190037 
## 
## Coefficients:
##             Estimate Std. Error t value Pr(>|t|)    
## (Intercept)  2.82550    0.01175 240.427   <2e-16 ***
## agejuv       0.08506    0.03716   2.289   0.0265 *  
## ---
## Signif. codes:  0 '***' 0.001 '**' 0.01 '*' 0.05 '.' 0.1 ' ' 1
## 
## Residual standard error: 0.07883 on 48 degrees of freedom
## Multiple R-squared:  0.0984, Adjusted R-squared:  0.07961 
## F-statistic: 5.238 on 1 and 48 DF,  p-value: 0.02654
```

```
plot_model(BodyMass_age_mod, type="emm", terms=c("age"), mdrt.values="meansd", ppd=TRUE, line.size=1,dot.size=5)
```

```
residuals_BodyMass_mod <- resid(BodyMass_mod)
hist(residuals_BodyMass_mod)
```

```
qqnorm(residuals_BodyMass_mod)
qqline(residuals_BodyMass_mod)
```

```
shapiro.test(residuals_BodyMass_mod)
```

```
## 
##  Shapiro-Wilk normality test
## 
## data:  residuals_BodyMass_mod
## W = 0.98006, p-value = 0.5544
```

```
#FFM as body mass
BodyMass_mod_FFM<-lm(logFFM ~ season+sex, data=Data_FFM)
summary(BodyMass_mod_FFM)
```

```
## 
## Call:
## lm(formula = logFFM ~ season + sex, data = Data_FFM)
## 
## Residuals:
##      Min       1Q   Median       3Q      Max 
## -0.16623 -0.04795  0.01610  0.04614  0.10653 
## 
## Coefficients:
##             Estimate Std. Error t value Pr(>|t|)    
## (Intercept) 2.744173   0.030381  90.325   <2e-16 ***
## seasonw     0.001898   0.045062   0.042    0.967    
## sexM        0.021881   0.047066   0.465    0.650    
## ---
## Signif. codes:  0 '***' 0.001 '**' 0.01 '*' 0.05 '.' 0.1 ' ' 1
## 
## Residual standard error: 0.07442 on 13 degrees of freedom
## Multiple R-squared:  0.02459,    Adjusted R-squared:  -0.1255 
## F-statistic: 0.1639 on 2 and 13 DF,  p-value: 0.8506
```

```
plot_model(BodyMass_mod_FFM, type="emm", terms=c("sex"), mdrt.values="meansd", ppd=TRUE, line.size=1,dot.size=5)
```

### 8 FMR ~ season

```
FMR_mod<-lm(resFMR ~ season, data=Data)
summary(FMR_mod)
```

```
## 
## Call:
## lm(formula = resFMR ~ season, data = Data)
## 
## Residuals:
##       16       29       42       45       53 
##  0.08795 -0.08795  0.17940 -0.01223 -0.16717 
## 
## Coefficients:
##             Estimate Std. Error t value Pr(>|t|)
## (Intercept)  0.01558    0.09174   0.170    0.876
## seasonw     -0.03894    0.14506  -0.268    0.806
## 
## Residual standard error: 0.1589 on 3 degrees of freedom
##   (51 observations deleted due to missingness)
## Multiple R-squared:  0.02346,    Adjusted R-squared:  -0.3021 
## F-statistic: 0.07206 on 1 and 3 DF,  p-value: 0.8058
```

```
plot_model(FMR_mod, type="emm", terms=c("season"), mdrt.values="meansd", ppd=TRUE, line.size=1,dot.size=5)
```

```
emmeans(FMR_mod, pairwise ~ season, nesting=NULL)
```

```
## $emmeans
##  season  emmean     SE df lower.CL upper.CL
##  b       0.0156 0.0917  3   -0.276    0.308
##  w      -0.0234 0.1124  3   -0.381    0.334
## 
## Confidence level used: 0.95 
## 
## $contrasts
##  contrast estimate    SE df t.ratio p.value
##  b - w      0.0389 0.145  3   0.268  0.8058
```

```
residuals_FMR_mod <- resid(FMR_mod)
hist(residuals_FMR_mod)
```

```
qqnorm(residuals_FMR_mod)
qqline(residuals_FMR_mod)
```

```
shapiro.test(residuals_FMR_mod)
```

```
## 
##  Shapiro-Wilk normality test
## 
## data:  residuals_FMR_mod
## W = 0.98243, p-value = 0.9472
```

```
#FFM

FMR_mod<-lm(resFMR_FFM ~ season, data=Data_FFM)
summary(FMR_mod)
```

```
## 
## Call:
## lm(formula = resFMR_FFM ~ season, data = Data_FFM)
## 
## Residuals:
##        7       10       12       14       16 
##  0.01721 -0.01721  0.17364  0.00128 -0.17492 
## 
## Coefficients:
##             Estimate Std. Error t value Pr(>|t|)
## (Intercept) -0.01933    0.08256  -0.234    0.830
## seasonw      0.04832    0.13053   0.370    0.736
## 
## Residual standard error: 0.143 on 3 degrees of freedom
##   (11 observations deleted due to missingness)
## Multiple R-squared:  0.04367,    Adjusted R-squared:  -0.2751 
## F-statistic: 0.137 on 1 and 3 DF,  p-value: 0.7359
```

```
plot_model(FMR_mod, type="emm", terms=c("season"), mdrt.values="meansd", ppd=TRUE, line.size=1,dot.size=5)
```

```
emmeans(FMR_mod, pairwise ~ season, nesting=NULL)
```

```
## $emmeans
##  season  emmean     SE df lower.CL upper.CL
##  b      -0.0193 0.0826  3   -0.282    0.243
##  w       0.0290 0.1011  3   -0.293    0.351
## 
## Confidence level used: 0.95 
## 
## $contrasts
##  contrast estimate    SE df t.ratio p.value
##  b - w     -0.0483 0.131  3  -0.370  0.7359
```

### 9 BMR (winter) ~ sex

#### 9.1 Mass-independent

```
BMR_w_mod<-lm(resBMR ~ sex, data=BMR_w)
plot_model(BMR_w_mod, type="emm", terms=c("sex"), mdrt.values="meansd", ppd=TRUE, line.size=1,dot.size=5)
```

```
emmeans(BMR_w_mod, pairwise ~ sex, nesting=NULL)
```

```
## $emmeans
##  sex   emmean     SE df lower.CL upper.CL
##  F    0.00895 0.0274 38  -0.0466   0.0645
##  M   -0.00989 0.0288 38  -0.0682   0.0485
## 
## Confidence level used: 0.95 
## 
## $contrasts
##  contrast estimate     SE df t.ratio p.value
##  F - M      0.0188 0.0398 38   0.473  0.6387
```

```
residuals_BMR_w_mod <- resid(BMR_w_mod)
hist(residuals_BMR_w_mod)
```

```
qqnorm(residuals_BMR_w_mod)
qqline(residuals_BMR_w_mod)
```

```
shapiro.test(residuals_BMR_w_mod)
```

```
## 
##  Shapiro-Wilk normality test
## 
## data:  residuals_BMR_w_mod
## W = 0.97685, p-value = 0.5742
```

#### 9.2 Whole-body

```
BMR_w_mod<-lm(logBMR ~ sex, data=BMR_w)
plot_model(BMR_w_mod, type="emm", terms=c("sex"), mdrt.values="meansd", ppd=TRUE, line.size=1,dot.size=5)
```

```
emmeans(BMR_w_mod, pairwise ~ sex, nesting=NULL)
```

```
## $emmeans
##  sex emmean     SE df lower.CL upper.CL
##  F    0.120 0.0302 38   0.0589    0.181
##  M    0.168 0.0318 38   0.1039    0.232
## 
## Confidence level used: 0.95 
## 
## $contrasts
##  contrast estimate     SE df t.ratio p.value
##  F - M     -0.0481 0.0438 38  -1.098  0.2791
```

```
residuals_BMR_w_mod <- resid(BMR_w_mod)
hist(residuals_BMR_w_mod)
```

```
qqnorm(residuals_BMR_w_mod)
qqline(residuals_BMR_w_mod)
```

```
shapiro.test(residuals_BMR_w_mod)
```

```
## 
##  Shapiro-Wilk normality test
## 
## data:  residuals_BMR_w_mod
## W = 0.9617, p-value = 0.1915
```

### 10 BMR ~ sex

#### 10.1 Mass-independent

```
BMR_w_mod<-lm(resBMR ~ sex, data=Data)
plot_model(BMR_w_mod, type="emm", terms=c("sex"), mdrt.values="meansd", ppd=TRUE, line.size=1,dot.size=5)
```

```
emmeans(BMR_w_mod, pairwise ~ sex, nesting=NULL)
```

```
## $emmeans
##  sex  emmean     SE df lower.CL upper.CL
##  F    0.0135 0.0248 48  -0.0364   0.0634
##  M   -0.0220 0.0317 48  -0.0857   0.0417
## 
## Confidence level used: 0.95 
## 
## $contrasts
##  contrast estimate     SE df t.ratio p.value
##  F - M      0.0355 0.0403 48   0.881  0.3827
```

```
residuals_BMR_w_mod <- resid(BMR_w_mod)
hist(residuals_BMR_w_mod)
```

```
qqnorm(residuals_BMR_w_mod)
qqline(residuals_BMR_w_mod)
```

```
shapiro.test(residuals_BMR_w_mod)
```

```
## 
##  Shapiro-Wilk normality test
## 
## data:  residuals_BMR_w_mod
## W = 0.98316, p-value = 0.6907
```

#### 10.2 Whole-body

```
BMR_w_mod<-lm(logBMR ~ sex, data=Data)
plot_model(BMR_w_mod, type="emm", terms=c("sex"), mdrt.values="meansd", ppd=TRUE, line.size=1,dot.size=5)
```

```
emmeans(BMR_w_mod, pairwise ~ sex, nesting=NULL)
```

```
## $emmeans
##  sex emmean     SE df lower.CL upper.CL
##  F    0.154 0.0264 48      0.1    0.207
##  M    0.168 0.0338 48      0.1    0.236
## 
## Confidence level used: 0.95 
## 
## $contrasts
##  contrast estimate     SE df t.ratio p.value
##  F - M     -0.0146 0.0429 48  -0.340  0.7350
```

### 11 Msum (winter) ~ sex

#### 11.1 Mass-independent

```
Msum_w_mod<-lm(resMsum ~ sex, data=Msum_w)
plot_model(Msum_w_mod, type="emm", terms=c("sex"), mdrt.values="meansd", ppd=TRUE, line.size=1,dot.size=5)
```

```
emmeans(Msum_w_mod, pairwise ~ sex, nesting=NULL)
```

```
## $emmeans
##  sex  emmean     SE df lower.CL upper.CL
##  F   -0.0206 0.0286 40  -0.0784   0.0373
##  M    0.0206 0.0286 40  -0.0373   0.0784
## 
## Confidence level used: 0.95 
## 
## $contrasts
##  contrast estimate     SE df t.ratio p.value
##  F - M     -0.0411 0.0405 40  -1.016  0.3159
```

```
residuals_Msum_w_mod <- resid(Msum_w_mod)
hist(residuals_Msum_w_mod)
```

```
qqnorm(residuals_Msum_w_mod)
qqline(residuals_Msum_w_mod)
```

```
shapiro.test(residuals_Msum_w_mod)
```

```
## 
##  Shapiro-Wilk normality test
## 
## data:  residuals_Msum_w_mod
## W = 0.96459, p-value = 0.215
```

#### 11.2 Whole-body

```
Msum_w_mod<-lm(logMsum ~ sex, data=Msum_w)
plot_model(Msum_w_mod, type="emm", terms=c("sex"), mdrt.values="meansd", ppd=TRUE, line.size=1,dot.size=5)
```

```
emmeans(Msum_w_mod, pairwise ~ sex, nesting=NULL)
```

```
## $emmeans
##  sex emmean     SE df lower.CL upper.CL
##  F     1.44 0.0291 40     1.38     1.50
##  M     1.56 0.0291 40     1.50     1.62
## 
## Confidence level used: 0.95 
## 
## $contrasts
##  contrast estimate     SE df t.ratio p.value
##  F - M      -0.125 0.0411 40  -3.048  0.0041
```

```
residuals_Msum_w_mod <- resid(Msum_w_mod)
hist(residuals_Msum_w_mod)
```

```
qqnorm(residuals_Msum_w_mod)
qqline(residuals_Msum_w_mod)
```

```
shapiro.test(residuals_Msum_w_mod)
```

```
## 
##  Shapiro-Wilk normality test
## 
## data:  residuals_Msum_w_mod
## W = 0.93093, p-value = 0.01392
```

### 12 BMR (winter) ~ age

#### 12.1 Mass-independent

```
BMR_w_mod2<-lm(resBMR ~ age, data=BMR_w)
plot_model(BMR_w_mod2, type="emm", terms=c("age"), mdrt.values="meansd", ppd=TRUE, line.size=1,dot.size=5)
```

```
emmeans(BMR_w_mod2, pairwise ~ age, nesting=NULL)
```

```
## $emmeans
##  age   emmean     SE df lower.CL upper.CL
##  ad  -0.00788 0.0210 38  -0.0504   0.0346
##  juv  0.05513 0.0555 38  -0.0573   0.1676
## 
## Confidence level used: 0.95 
## 
## $contrasts
##  contrast estimate     SE df t.ratio p.value
##  ad - juv   -0.063 0.0594 38  -1.061  0.2953
```

```
residuals_BMR_w_mod2 <- resid(BMR_w_mod2)
hist(residuals_BMR_w_mod2)
```

```
qqnorm(residuals_BMR_w_mod2)
qqline(residuals_BMR_w_mod2)
```

```
shapiro.test(residuals_BMR_w_mod2)
```

```
## 
##  Shapiro-Wilk normality test
## 
## data:  residuals_BMR_w_mod2
## W = 0.9844, p-value = 0.8454
```

#### 12.2 Whole-body

```
BMR_w_mod2<-lm(logBMR ~ age, data=BMR_w)
plot_model(BMR_w_mod2, type="emm", terms=c("age"), mdrt.values="meansd", ppd=TRUE, line.size=1,dot.size=5)
```

```
emmeans(BMR_w_mod2, pairwise ~ age, nesting=NULL)
```

```
## $emmeans
##  age emmean     SE df lower.CL upper.CL
##  ad   0.128 0.0227 38   0.0817    0.174
##  juv  0.249 0.0601 38   0.1274    0.371
## 
## Confidence level used: 0.95 
## 
## $contrasts
##  contrast estimate     SE df t.ratio p.value
##  ad - juv   -0.121 0.0643 38  -1.889  0.0666
```

```
residuals_BMR_w_mod2 <- resid(BMR_w_mod2)
hist(residuals_BMR_w_mod2)
```

```
qqnorm(residuals_BMR_w_mod2)
qqline(residuals_BMR_w_mod2)
```

```
shapiro.test(residuals_BMR_w_mod2)
```

```
## 
##  Shapiro-Wilk normality test
## 
## data:  residuals_BMR_w_mod2
## W = 0.9737, p-value = 0.4674
```

```
ggplot(BMR_w, aes(age,resBMR))+
geom_violin(position="dodge",aes(fill=age),show.legend = FALSE)+labs(x="",y="mass-independent BMR")+ scale_x_discrete(limits = c("ad", "juv")) +
  scale_fill_manual(values=wes_palette(n=3, "GrandBudapest1"))+
  ggtitle("BMR between ad and juv in winter")+
  geom_dotplot(binaxis='y', stackdir='center', dotsize=0.5)+
  theme_light()+
  theme(panel.grid.minor = element_blank())+
  theme(strip.background = element_rect(fill = "white"))+
  theme(strip.text = element_text(colour = 'black',face="bold"))
```

### 13 Female BMR ~ season

#### 13.1 Mass-independent

```
BMR_f_mod <- lm(resBMR ~ season, data=BMR_f)
summary(BMR_f_mod)
```

```
## 
## Call:
## lm(formula = resBMR ~ season, data = BMR_f)
## 
## Residuals:
##      Min       1Q   Median       3Q      Max 
## -0.28518 -0.08568  0.01004  0.06628  0.35059 
## 
## Coefficients:
##             Estimate Std. Error t value Pr(>|t|)  
## (Intercept)  0.07256    0.04246   1.709   0.0939 .
## seasonw     -0.09070    0.04747  -1.911   0.0620 .
## ---
## Signif. codes:  0 '***' 0.001 '**' 0.01 '*' 0.05 '.' 0.1 ' ' 1
## 
## Residual standard error: 0.1343 on 48 degrees of freedom
## Multiple R-squared:  0.07068,    Adjusted R-squared:  0.05131 
## F-statistic:  3.65 on 1 and 48 DF,  p-value: 0.06204
```

```
plot_model(BMR_f_mod, type="emm", terms=c("season"), mdrt.values="meansd", ppd=TRUE, line.size=1,dot.size=5)
```

```
emmeans(BMR_f_mod, pairwise ~ season, nesting=NULL)
```

```
## $emmeans
##  season  emmean     SE df lower.CL upper.CL
##  b       0.0726 0.0425 48  -0.0128   0.1579
##  w      -0.0181 0.0212 48  -0.0608   0.0245
## 
## Confidence level used: 0.95 
## 
## $contrasts
##  contrast estimate     SE df t.ratio p.value
##  b - w      0.0907 0.0475 48   1.911  0.0620
```

```
residuals_BMR_f_mod <- resid(BMR_f_mod)
hist(residuals_BMR_f_mod)
```

```
qqnorm(residuals_BMR_f_mod)
qqline(residuals_BMR_f_mod)
```

```
shapiro.test(residuals_BMR_f_mod)
```

```
## 
##  Shapiro-Wilk normality test
## 
## data:  residuals_BMR_f_mod
## W = 0.98581, p-value = 0.8058
```

#### 13.2 Whole-body

```
BMR_f_mod <- lm(logBMR ~ season, data=BMR_f)
summary(BMR_f_mod)
```

```
## 
## Call:
## lm(formula = logBMR ~ season, data = BMR_f)
## 
## Residuals:
##      Min       1Q   Median       3Q      Max 
## -0.29369 -0.08381  0.00907  0.06771  0.44503 
## 
## Coefficients:
##             Estimate Std. Error t value Pr(>|t|)    
## (Intercept)  0.22402    0.04540   4.935 1.01e-05 ***
## seasonw     -0.08110    0.05075  -1.598    0.117    
## ---
## Signif. codes:  0 '***' 0.001 '**' 0.01 '*' 0.05 '.' 0.1 ' ' 1
## 
## Residual standard error: 0.1436 on 48 degrees of freedom
## Multiple R-squared:  0.0505, Adjusted R-squared:  0.03072 
## F-statistic: 2.553 on 1 and 48 DF,  p-value: 0.1166
```

```
plot_model(BMR_f_mod, type="emm", terms=c("season"), mdrt.values="meansd", ppd=TRUE, line.size=1,dot.size=5)
```

```
emmeans(BMR_f_mod, pairwise ~ season, nesting=NULL)
```

```
## $emmeans
##  season emmean     SE df lower.CL upper.CL
##  b       0.224 0.0454 48   0.1327    0.315
##  w       0.143 0.0227 48   0.0973    0.189
## 
## Confidence level used: 0.95 
## 
## $contrasts
##  contrast estimate     SE df t.ratio p.value
##  b - w      0.0811 0.0508 48   1.598  0.1166
```

```
residuals_BMR_f_mod <- resid(BMR_f_mod)
hist(residuals_BMR_f_mod)
```

```
qqnorm(residuals_BMR_f_mod)
qqline(residuals_BMR_f_mod)
```

```
shapiro.test(residuals_BMR_f_mod)
```

```
## 
##  Shapiro-Wilk normality test
## 
## data:  residuals_BMR_f_mod
## W = 0.97635, p-value = 0.4104
```

```
ggplot(BMR_f, aes(season,resBMR))+
geom_violin(position="dodge",aes(fill=season),show.legend = FALSE)+labs(x="",y="mass-independent BMR")+ scale_x_discrete(limits = c("b", "w")) +
  scale_fill_manual(values=wes_palette(n=3, "GrandBudapest1"))+
  ggtitle("F BMR between seasons")+
  geom_dotplot(binaxis='y', stackdir='center', dotsize=0.5)+
  theme_light()+
  theme(panel.grid.minor = element_blank())+
  theme(strip.background = element_rect(fill = "white"))+
  theme(strip.text = element_text(colour = 'black',face="bold"))
```

### 14 Msum (winter) ~ age

#### 14.1 Mass-independent

```
Msum_w_mod2<-lm(resMsum ~ age, data=Data_NoOL)
plot_model(Msum_w_mod2, type="emm", terms=c("age"), mdrt.values="meansd", ppd=TRUE, line.size=1,dot.size=5)
```

```
emmeans(Msum_w_mod2, pairwise ~ age, nesting=NULL)
```

```
## $emmeans
##  age  emmean     SE df lower.CL upper.CL
##  ad  -0.0131 0.0158 38  -0.0450   0.0188
##  juv  0.0990 0.0417 38   0.0146   0.1834
## 
## Confidence level used: 0.95 
## 
## $contrasts
##  contrast estimate     SE df t.ratio p.value
##  ad - juv   -0.112 0.0446 38  -2.514  0.0163
```

```
residuals_Msum_w_mod2 <- resid(Msum_w_mod2)
hist(residuals_Msum_w_mod2)
```

```
qqnorm(residuals_Msum_w_mod2)
qqline(residuals_Msum_w_mod2)
```

```
shapiro.test(residuals_Msum_w_mod2)
```

```
## 
##  Shapiro-Wilk normality test
## 
## data:  residuals_Msum_w_mod2
## W = 0.98523, p-value = 0.8713
```

#### 14.2 Whole-body

```
Msum_w_mod2<-lm(logMsum ~ age, data=Data_NoOL)
plot_model(Msum_w_mod2, type="emm", terms=c("age"), mdrt.values="meansd", ppd=TRUE, line.size=1,dot.size=5)
```

```
emmeans(Msum_w_mod2, pairwise ~ age, nesting=NULL)
```

```
## $emmeans
##  age emmean     SE df lower.CL upper.CL
##  ad    1.48 0.0148 38     1.45     1.51
##  juv   1.65 0.0391 38     1.57     1.73
## 
## Confidence level used: 0.95 
## 
## $contrasts
##  contrast estimate     SE df t.ratio p.value
##  ad - juv   -0.169 0.0418 38  -4.054  0.0002
```

```
residuals_Msum_w_mod2 <- resid(Msum_w_mod2)
hist(residuals_Msum_w_mod2)
```

```
qqnorm(residuals_Msum_w_mod2)
qqline(residuals_Msum_w_mod2)
```

```
shapiro.test(residuals_Msum_w_mod2)
```

```
## 
##  Shapiro-Wilk normality test
## 
## data:  residuals_Msum_w_mod2
## W = 0.98025, p-value = 0.6988
```

```
Msum_w_mod2_test<-lm(logmassMsum_1 ~ age, data=Data_NoOL)
plot_model(Msum_w_mod2_test, type="emm", terms=c("age"), mdrt.values="meansd", ppd=TRUE, line.size=1,dot.size=5)
```

```
emmeans(Msum_w_mod2_test, pairwise ~ age, nesting=NULL)
```

```
## $emmeans
##  age emmean     SE df lower.CL upper.CL
##  ad    2.79 0.0123 38     2.77     2.82
##  juv   2.86 0.0324 38     2.80     2.93
## 
## Confidence level used: 0.95 
## 
## $contrasts
##  contrast estimate     SE df t.ratio p.value
##  ad - juv  -0.0684 0.0347 38  -1.973  0.0558
```

```
residuals_Msum_w_mod2_test <- resid(Msum_w_mod2_test)
hist(residuals_Msum_w_mod2_test)
```

```
qqnorm(residuals_Msum_w_mod2_test)
qqline(residuals_Msum_w_mod2_test)
```

```
shapiro.test(residuals_Msum_w_mod2_test)
```

```
## 
##  Shapiro-Wilk normality test
## 
## data:  residuals_Msum_w_mod2_test
## W = 0.98476, p-value = 0.8569
```

```
ggplot(Data_NoOL, aes(age,resMsum))+
geom_violin(position="dodge",aes(fill=age),show.legend = FALSE)+labs(x="",y="mass-independent Msum")+ scale_x_discrete(limits = c("ad", "juv")) +
  scale_fill_manual(values=wes_palette(n=3, "GrandBudapest1"))+
  ggtitle("Msum between ad and juv in winter")+
  geom_dotplot(binaxis='y', stackdir='center', dotsize=0.5)+
  theme_light()+
  theme(panel.grid.minor = element_blank())+
  theme(strip.background = element_rect(fill = "white"))+
  theme(strip.text = element_text(colour = 'black',face="bold"))
```

### 15 BMR ~ season

#### 15.1 Mass-independent

```
BMR_mod<-lm(resBMR ~ season, data=Data_BMR)
summary(BMR_mod)
```

```
## 
## Call:
## lm(formula = resBMR ~ season, data = Data_BMR)
## 
## Residuals:
##      Min       1Q   Median       3Q      Max 
## -0.28518 -0.08568  0.01004  0.06628  0.35059 
## 
## Coefficients:
##             Estimate Std. Error t value Pr(>|t|)  
## (Intercept)  0.07256    0.04246   1.709   0.0939 .
## seasonw     -0.09070    0.04747  -1.911   0.0620 .
## ---
## Signif. codes:  0 '***' 0.001 '**' 0.01 '*' 0.05 '.' 0.1 ' ' 1
## 
## Residual standard error: 0.1343 on 48 degrees of freedom
## Multiple R-squared:  0.07068,    Adjusted R-squared:  0.05131 
## F-statistic:  3.65 on 1 and 48 DF,  p-value: 0.06204
```

```
plot_model(BMR_mod, type="emm", terms=c("season"), mdrt.values="meansd", ppd=TRUE, line.size=1,dot.size=5)
```

```
emmeans(BMR_mod, pairwise ~ season, nesting=NULL)
```

```
## $emmeans
##  season  emmean     SE df lower.CL upper.CL
##  b       0.0726 0.0425 48  -0.0128   0.1579
##  w      -0.0181 0.0212 48  -0.0608   0.0245
## 
## Confidence level used: 0.95 
## 
## $contrasts
##  contrast estimate     SE df t.ratio p.value
##  b - w      0.0907 0.0475 48   1.911  0.0620
```

```
residuals_BMR_mod <- resid(BMR_mod)
hist(residuals_BMR_mod)
```

```
qqnorm(residuals_BMR_mod)
qqline(residuals_BMR_mod)
```

```
shapiro.test(residuals_BMR_mod)
```

```
## 
##  Shapiro-Wilk normality test
## 
## data:  residuals_BMR_mod
## W = 0.98581, p-value = 0.8058
```

```
#FFM

BMR_mod_FFM <- lm(resBMR_FFM ~ season+sex, data=Data_FFM)
summary(BMR_mod_FFM)
```

```
## 
## Call:
## lm(formula = resBMR_FFM ~ season + sex, data = Data_FFM)
## 
## Residuals:
##      Min       1Q   Median       3Q      Max 
## -0.31889 -0.06418  0.02530  0.08954  0.21435 
## 
## Coefficients:
##             Estimate Std. Error t value Pr(>|t|)
## (Intercept)  0.04720    0.06519   0.724    0.483
## seasonw     -0.11731    0.09219  -1.272    0.227
## sexM         0.09302    0.09219   1.009    0.333
## 
## Residual standard error: 0.1458 on 12 degrees of freedom
##   (1 observation deleted due to missingness)
## Multiple R-squared:  0.1307, Adjusted R-squared:  -0.01418 
## F-statistic: 0.9021 on 2 and 12 DF,  p-value: 0.4315
```

```
Msum_mod_FFM <- lm(resMsum_FFM ~ season+sex, data=Data_FFM)
summary(Msum_mod_FFM)
```

```
## 
## Call:
## lm(formula = resMsum_FFM ~ season + sex, data = Data_FFM)
## 
## Residuals:
##      Min       1Q   Median       3Q      Max 
## -0.14395 -0.09209  0.00000  0.01682  0.29410 
## 
## Coefficients:
##             Estimate Std. Error t value Pr(>|t|)
## (Intercept) -0.19277    0.13885  -1.388    0.202
## seasonw      0.17403    0.15210   1.144    0.286
## sexM         0.07604    0.08782   0.866    0.412
## 
## Residual standard error: 0.1388 on 8 degrees of freedom
##   (5 observations deleted due to missingness)
## Multiple R-squared:  0.264,  Adjusted R-squared:  0.08004 
## F-statistic: 1.435 on 2 and 8 DF,  p-value: 0.2934
```

#### 15.2 Whole-body

```
BMR_mod<-lm(logBMR ~ season, data=Data_BMR)
summary(BMR_mod)
```

```
## 
## Call:
## lm(formula = logBMR ~ season, data = Data_BMR)
## 
## Residuals:
##      Min       1Q   Median       3Q      Max 
## -0.29369 -0.08381  0.00907  0.06771  0.44503 
## 
## Coefficients:
##             Estimate Std. Error t value Pr(>|t|)    
## (Intercept)  0.22402    0.04540   4.935 1.01e-05 ***
## seasonw     -0.08110    0.05075  -1.598    0.117    
## ---
## Signif. codes:  0 '***' 0.001 '**' 0.01 '*' 0.05 '.' 0.1 ' ' 1
## 
## Residual standard error: 0.1436 on 48 degrees of freedom
## Multiple R-squared:  0.0505, Adjusted R-squared:  0.03072 
## F-statistic: 2.553 on 1 and 48 DF,  p-value: 0.1166
```

```
plot_model(BMR_mod, type="emm", terms=c("season"), mdrt.values="meansd", ppd=TRUE, line.size=1,dot.size=5)
```

```
emmeans(BMR_mod, pairwise ~ season, nesting=NULL)
```

```
## $emmeans
##  season emmean     SE df lower.CL upper.CL
##  b       0.224 0.0454 48   0.1327    0.315
##  w       0.143 0.0227 48   0.0973    0.189
## 
## Confidence level used: 0.95 
## 
## $contrasts
##  contrast estimate     SE df t.ratio p.value
##  b - w      0.0811 0.0508 48   1.598  0.1166
```

```
residuals_BMR_mod <- resid(BMR_mod)
hist(residuals_BMR_mod)
```

```
qqnorm(residuals_BMR_mod)
qqline(residuals_BMR_mod)
```

```
shapiro.test(residuals_BMR_mod)
```

```
## 
##  Shapiro-Wilk normality test
## 
## data:  residuals_BMR_mod
## W = 0.97635, p-value = 0.4104
```

```
ggplot(Data_BMR, aes(season,resBMR))+
geom_violin(position="dodge",aes(fill=season),show.legend = FALSE)+labs(x="",y="mass-independent BMR")+ scale_x_discrete(limits = c("b", "w")) +
  scale_fill_manual(values=wes_palette(n=3, "GrandBudapest1"))+
  ggtitle("BMR between seasons")+
  geom_dotplot(binaxis='y', stackdir='center', dotsize=0.5)+
  theme_light()+
  theme(panel.grid.minor = element_blank())+
  theme(strip.background = element_rect(fill = "white"))+
  theme(strip.text = element_text(colour = 'black',face="bold"))
```

### 16 Msum ~ season

#### 16.1 Whole-body

```
Msum_mod1<-lm(logMsum ~ season, data=Data_NoOL)
summary(Msum_mod1)
```

```
## 
## Call:
## lm(formula = logMsum ~ season, data = Data_NoOL)
## 
## Residuals:
##       Min        1Q    Median        3Q       Max 
## -0.199887 -0.075968 -0.000204  0.057571  0.228038 
## 
## Coefficients:
##             Estimate Std. Error t value Pr(>|t|)    
## (Intercept)  1.44993    0.04164  34.817   <2e-16 ***
## seasonw      0.06245    0.04517   1.383    0.175    
## ---
## Signif. codes:  0 '***' 0.001 '**' 0.01 '*' 0.05 '.' 0.1 ' ' 1
## 
## Residual standard error: 0.102 on 38 degrees of freedom
## Multiple R-squared:  0.04789,    Adjusted R-squared:  0.02284 
## F-statistic: 1.911 on 1 and 38 DF,  p-value: 0.1749
```

```
plot_model(Msum_mod1, type="emm", terms=c("season"), mdrt.values="meansd", ppd=TRUE, line.size=1,dot.size=5)
```

```
emmeans(Msum_mod1, pairwise ~ season, nesting=NULL)
```

```
## $emmeans
##  season emmean     SE df lower.CL upper.CL
##  b        1.45 0.0416 38     1.37     1.53
##  w        1.51 0.0175 38     1.48     1.55
## 
## Confidence level used: 0.95 
## 
## $contrasts
##  contrast estimate     SE df t.ratio p.value
##  b - w     -0.0624 0.0452 38  -1.383  0.1749
```

```
residuals_Msum_mod1 <- resid(Msum_mod1)
hist(residuals_Msum_mod1)
```

```
qqnorm(residuals_Msum_mod1)
qqline(residuals_Msum_mod1)
```

```
shapiro.test(residuals_Msum_mod1)
```

```
## 
##  Shapiro-Wilk normality test
## 
## data:  residuals_Msum_mod1
## W = 0.99056, p-value = 0.981
```

#### 16.2 Mass-independent

```
Msum_mod<-lm(resMsum ~ season, data=Data_NoOL)
summary(Msum_mod)
```

```
## 
## Call:
## lm(formula = resMsum ~ season, data = Data_NoOL)
## 
## Residuals:
##       Min        1Q    Median        3Q       Max 
## -0.192672 -0.064601 -0.008924  0.047661  0.183411 
## 
## Coefficients:
##             Estimate Std. Error t value Pr(>|t|)   
## (Intercept) -0.09885    0.03717  -2.659   0.0114 * 
## seasonw      0.11735    0.04032   2.911   0.0060 **
## ---
## Signif. codes:  0 '***' 0.001 '**' 0.01 '*' 0.05 '.' 0.1 ' ' 1
## 
## Residual standard error: 0.09105 on 38 degrees of freedom
## Multiple R-squared:  0.1823, Adjusted R-squared:  0.1608 
## F-statistic: 8.472 on 1 and 38 DF,  p-value: 0.006003
```

```
plot_model(Msum_mod, type="emm", terms=c("season"), mdrt.values="meansd", ppd=TRUE, line.size=1,dot.size=5)
```

```
emmeans(Msum_mod, pairwise ~ season, nesting=NULL)
```

```
## $emmeans
##  season  emmean     SE df lower.CL upper.CL
##  b      -0.0988 0.0372 38  -0.1741  -0.0236
##  w       0.0185 0.0156 38  -0.0131   0.0501
## 
## Confidence level used: 0.95 
## 
## $contrasts
##  contrast estimate     SE df t.ratio p.value
##  b - w      -0.117 0.0403 38  -2.911  0.0060
```

```
residuals_Msum_mod <- resid(Msum_mod)
hist(residuals_Msum_mod)
```

```
qqnorm(residuals_Msum_mod)
qqline(residuals_Msum_mod)
```

```
shapiro.test(residuals_Msum_mod)
```

```
## 
##  Shapiro-Wilk normality test
## 
## data:  residuals_Msum_mod
## W = 0.97614, p-value = 0.5492
```

```
Msum_mod<-lm(resMsum_FFM ~ season, data=Data_FFM)
summary(Msum_mod)
```

```
## 
## Call:
## lm(formula = resMsum_FFM ~ season, data = Data_FFM)
## 
## Residuals:
##      Min       1Q   Median       3Q      Max 
## -0.18197 -0.05407 -0.03662  0.02224  0.33212 
## 
## Coefficients:
##             Estimate Std. Error t value Pr(>|t|)
## (Intercept)  -0.1928     0.1369  -1.408    0.193
## seasonw       0.2120     0.1436   1.477    0.174
## 
## Residual standard error: 0.1369 on 9 degrees of freedom
##   (5 observations deleted due to missingness)
## Multiple R-squared:  0.1951, Adjusted R-squared:  0.1056 
## F-statistic: 2.181 on 1 and 9 DF,  p-value: 0.1738
```

```
plot_model(Msum_mod, type="emm", terms=c("season"), mdrt.values="meansd", ppd=TRUE, line.size=1,dot.size=5)
```

```
emmeans(Msum_mod, pairwise ~ season, nesting=NULL)
```

```
## $emmeans
##  season  emmean     SE df lower.CL upper.CL
##  b      -0.1928 0.1369  9  -0.5025    0.117
##  w       0.0193 0.0433  9  -0.0787    0.117
## 
## Confidence level used: 0.95 
## 
## $contrasts
##  contrast estimate    SE df t.ratio p.value
##  b - w      -0.212 0.144  9  -1.477  0.1738
```

```
ggplot(Data_NoOL, aes(season,resMsum))+
geom_violin(position="dodge",aes(fill=season),show.legend = FALSE)+labs(x="",y="mass-independent Msum")+ scale_x_discrete(limits = c("b", "w")) +
  scale_fill_manual(values=wes_palette(n=3, "GrandBudapest1"))+
  ggtitle("Msum between seasons")+
  geom_dotplot(binaxis='y', stackdir='center', dotsize=0.5)+
  theme_light()+
  theme(panel.grid.minor = element_blank())+
  theme(strip.background = element_rect(fill = "white"))+
  theme(strip.text = element_text(colour = 'black',face="bold"))
```

### 17 Cold tolerance ~ season

```
Msum_mod3<-lm(T_end ~ massMsum_1 + season, data=Data_NoOL)
summary(Msum_mod3)
```

```
## 
## Call:
## lm(formula = T_end ~ massMsum_1 + season, data = Data_NoOL)
## 
## Residuals:
##     Min      1Q  Median      3Q     Max 
## -2.3022 -1.1456 -0.1695  1.0775  3.7402 
## 
## Coefficients:
##             Estimate Std. Error t value Pr(>|t|)   
## (Intercept)  -8.1371     4.0864  -1.991   0.0550 . 
## massMsum_1   -0.4788     0.2313  -2.070   0.0466 * 
## seasonw      -2.3679     0.7249  -3.267   0.0026 **
## ---
## Signif. codes:  0 '***' 0.001 '**' 0.01 '*' 0.05 '.' 0.1 ' ' 1
## 
## Residual standard error: 1.506 on 32 degrees of freedom
##   (5 observations deleted due to missingness)
## Multiple R-squared:  0.2656, Adjusted R-squared:  0.2197 
## F-statistic: 5.788 on 2 and 32 DF,  p-value: 0.007153
```

```
plot_model(Msum_mod3, type="emm", terms=c("season"), mdrt.values="meansd", ppd=TRUE, line.size=1,dot.size=5)
```

```
emmeans(Msum_mod3, pairwise ~ season, nesting=NULL)
```

```
## $emmeans
##  season emmean    SE df lower.CL upper.CL
##  b       -16.0 0.652 32    -17.4    -14.7
##  w       -18.4 0.283 32    -19.0    -17.8
## 
## Confidence level used: 0.95 
## 
## $contrasts
##  contrast estimate    SE df t.ratio p.value
##  b - w        2.37 0.725 32   3.267  0.0026
```

```
residuals_Msum_mod3 <- resid(Msum_mod3)
hist(residuals_Msum_mod3)
```

```
qqnorm(residuals_Msum_mod3)
qqline(residuals_Msum_mod3)
```

```
shapiro.test(residuals_Msum_mod3)
```

```
## 
##  Shapiro-Wilk normality test
## 
## data:  residuals_Msum_mod3
## W = 0.96781, p-value = 0.386
```

### 18 Cold tolerance ~ Msum

#### 18.1 Mass-independent

```
TEst_CT_Msum_FFM<-lm(resMsum ~ massMsum_1 + season, data=Data_NoOL)
summary(Msum_mod3)
```

```
## 
## Call:
## lm(formula = T_end ~ massMsum_1 + season, data = Data_NoOL)
## 
## Residuals:
##     Min      1Q  Median      3Q     Max 
## -2.3022 -1.1456 -0.1695  1.0775  3.7402 
## 
## Coefficients:
##             Estimate Std. Error t value Pr(>|t|)   
## (Intercept)  -8.1371     4.0864  -1.991   0.0550 . 
## massMsum_1   -0.4788     0.2313  -2.070   0.0466 * 
## seasonw      -2.3679     0.7249  -3.267   0.0026 **
## ---
## Signif. codes:  0 '***' 0.001 '**' 0.01 '*' 0.05 '.' 0.1 ' ' 1
## 
## Residual standard error: 1.506 on 32 degrees of freedom
##   (5 observations deleted due to missingness)
## Multiple R-squared:  0.2656, Adjusted R-squared:  0.2197 
## F-statistic: 5.788 on 2 and 32 DF,  p-value: 0.007153
```

```
plot_model(Msum_mod3, type="emm", terms=c("season"), mdrt.values="meansd", ppd=TRUE, line.size=1,dot.size=5)
```

```
emmeans(Msum_mod3, pairwise ~ season, nesting=NULL)
```

```
## $emmeans
##  season emmean    SE df lower.CL upper.CL
##  b       -16.0 0.652 32    -17.4    -14.7
##  w       -18.4 0.283 32    -19.0    -17.8
## 
## Confidence level used: 0.95 
## 
## $contrasts
##  contrast estimate    SE df t.ratio p.value
##  b - w        2.37 0.725 32   3.267  0.0026
```

```
residuals_TEst_CT_Msum_FFM <- resid(TEst_CT_Msum_FFM)
hist(residuals_TEst_CT_Msum_FFM)
```

```
qqnorm(residuals_TEst_CT_Msum_FFM)
qqline(residuals_TEst_CT_Msum_FFM)
```

```
shapiro.test(residuals_TEst_CT_Msum_FFM)
```

```
## 
##  Shapiro-Wilk normality test
## 
## data:  residuals_TEst_CT_Msum_FFM
## W = 0.97279, p-value = 0.4391
```

```
T_cold_Msum<-ggscatter(Data_Msum, y = "T_end", x = "resMsum", 
          add = "reg.line", conf.int = TRUE, 
          cor.coef = TRUE, cor.method = "pearson",
          ylab = "Cold tolerance [°C]", xlab = "mass-independent Msum")
T_cold_Msum
```

```
#FFM

T_cold_Msum<-ggscatter(Data_FFM, y = "resMsum_FFM", x = "T_end", 
          add = "reg.line", conf.int = TRUE, 
          cor.coef = TRUE, cor.method = "pearson",
          ylab = "mass-independent Msum", xlab = "Cold tolerance [°C]")
T_cold_Msum
```

```
T_cold_Msum<-ggscatter(Data_FFM, y = "resMsum", x = "T_end", 
          add = "reg.line", conf.int = TRUE, 
          cor.coef = TRUE, cor.method = "pearson",
          ylab = "mass-independent Msum", xlab = "Cold tolerance [°C]")
T_cold_Msum
```

```
T_cold_Msum_FFM<-lm(resMsum_FFM ~ T_end, Data_FFM)
summary(T_cold_Msum_FFM)
```

```
## 
## Call:
## lm(formula = resMsum_FFM ~ T_end, data = Data_FFM)
## 
## Residuals:
##       Min        1Q    Median        3Q       Max 
## -0.046983 -0.032832 -0.008517  0.012413  0.084508 
## 
## Coefficients:
##             Estimate Std. Error t value Pr(>|t|)   
## (Intercept) -0.86758    0.20203  -4.294  0.00359 **
## T_end       -0.04423    0.01091  -4.055  0.00484 **
## ---
## Signif. codes:  0 '***' 0.001 '**' 0.01 '*' 0.05 '.' 0.1 ' ' 1
## 
## Residual standard error: 0.04689 on 7 degrees of freedom
##   (7 observations deleted due to missingness)
## Multiple R-squared:  0.7014, Adjusted R-squared:  0.6587 
## F-statistic: 16.44 on 1 and 7 DF,  p-value: 0.004842
```

```
AIC(T_cold_Msum_FFM)
```

```
## [1] -25.8
```

```
T_cold_Msum<-lm(resMsum ~ T_end, Data_FFM)
summary(T_cold_Msum)
```

```
## 
## Call:
## lm(formula = resMsum ~ T_end, data = Data_FFM)
## 
## Residuals:
##       Min        1Q    Median        3Q       Max 
## -0.043728 -0.030605  0.003915  0.019638  0.055684 
## 
## Coefficients:
##             Estimate Std. Error t value Pr(>|t|)    
## (Intercept) -0.95009    0.16800  -5.655 0.000770 ***
## T_end       -0.05106    0.00907  -5.629 0.000791 ***
## ---
## Signif. codes:  0 '***' 0.001 '**' 0.01 '*' 0.05 '.' 0.1 ' ' 1
## 
## Residual standard error: 0.03899 on 7 degrees of freedom
##   (7 observations deleted due to missingness)
## Multiple R-squared:  0.8191, Adjusted R-squared:  0.7932 
## F-statistic: 31.69 on 1 and 7 DF,  p-value: 0.0007913
```

```
AIC(T_cold_Msum)
```

```
## [1] -29.12031
```

#### 18.2 Whole-body

```
T_cold_Msum<-ggscatter(Data_Msum, y = "T_end", x = "logMsum", 
          add = "reg.line", conf.int = TRUE, 
          cor.coef = TRUE, cor.method = "pearson",
          ylab = "Cold tolerance [°C]", xlab = "whole-body Msum [ml/min]")
T_cold_Msum
```

### 19 Figures

```
annotate_figure(Plot,
               top = text_grob("Aerobic Capacity Model", color = "black", face = "bold", size = 28))
```

```
ggsave("ACM.png", Plot, width = 18, height = 11, dpi = 600)
```

```
dat_text_1 <- data.frame(label = c("                                                                         ns"),size=3)
plot5<-ggplot(Data, aes(season,logBMR))+
geom_violin(position="dodge",aes(fill=season),show.legend = FALSE)+
  geom_text(data=dat_text_1,
            mapping = aes(x="w",y = 0.66, label = label, vjust=0))+
  annotate("segment", x = "w", xend = "b", y = 0.65, yend = 0.65,
  colour = "black", size=1)+
  labs(x="",y="log [BMR], ml/min")+
  scale_x_discrete(limits = c("w", "b"))+
  scale_fill_manual(values=wes_palette(n=3, "GrandBudapest1"))+
  geom_dotplot(binaxis='y', stackdir='center', dotsize=0.75)+
  theme(strip.background = element_rect(fill = "white"), axis.title=element_text(size=15))+
  theme(strip.text = element_text(colour = 'black'))+
  theme_light()+
  theme(panel.grid.major = element_blank(), panel.grid.minor = element_blank(),
panel.background = element_blank())
plot5
```

```
dat_text_2 <- data.frame(label = c("                                                                         ns"),size=3)
plot6<-ggplot(Data_NoOL, aes(season,logMsum))+
geom_violin(position="dodge",aes(fill=season),show.legend = FALSE)+
  geom_text(data=dat_text_2,
            mapping = aes(x="w",y = 1.8, label = label, vjust=0))+
  annotate("segment", x = "w", xend = "b", y = 1.79, yend = 1.79,
  colour = "black", size=1)+
  labs(x="",y="log [Msum], ml/min")+
  scale_x_discrete(limits = c("w", "b"))+
  scale_fill_manual(values=wes_palette(n=3, "GrandBudapest1"))+
  geom_dotplot(binaxis='y', stackdir='center', dotsize=0.75)+
  theme(strip.background = element_rect(fill = "white"), axis.title=element_text(size=15))+
  theme(strip.text = element_text(colour = 'black'))+
  theme_light()+
  theme(panel.grid.major = element_blank(), panel.grid.minor = element_blank(),
panel.background = element_blank())
plot6
```

```
dat_text_3 <- data.frame(label = c("                                                                         ns"),size=3)
plot7<-ggplot(Data, aes(season,logFMR))+
geom_violin(position="dodge",aes(fill=season),show.legend = FALSE)+
  geom_text(data=dat_text_2,
            mapping = aes(x="w",y = 4.62, label = label, vjust=0))+
  annotate("segment", x = "w", xend = "b", y = 4.6, yend = 4.6,
  colour = "black", size=1)+
  labs(x="",y="log [FMR], kJ/day")+
  scale_x_discrete(limits = c("w", "b"))+
  scale_fill_manual(values=wes_palette(n=3, "GrandBudapest1"))+
  geom_dotplot(binaxis='y', stackdir='center', dotsize=0.75)+
  theme(strip.background = element_rect(fill = "white"), axis.title=element_text(size=15))+
  theme(strip.text = element_text(colour = 'black'))+
  theme_light()+
  theme(panel.grid.major = element_blank(), panel.grid.minor = element_blank(),
panel.background = element_blank())
plot7
```

### 20 Thermoneutral zone

```
Plot2<-ggplot(TNZ, aes(T, VO2_mean))+ 
  geom_point(size = 3)+
  geom_errorbar(aes(ymin=VO2_mean-VO2_SD, ymax=VO2_mean+VO2_SD), width =1, size=1)+
  labs(x="T, °C", y=expression(VO[2]~","~ml/min))+
  ylim(0.85,1.5)+
  theme(panel.grid.major = element_blank(), panel.grid.minor = element_blank())+
  theme_classic(base_size = 15)
Plot2
```

```
ggsave("TNZ.png", Plot2, width = 10, height = 8, dpi = 600)
```

### 21 Extra analysis

```
ggscatter(BMR_b, x = "resBMR", y = "chicks", 
          add = "reg.line", conf.int = TRUE, 
          cor.coef = TRUE, cor.method = "pearson",
          xlab = "mass-independent BMR", ylab = "Number of chicks")
```

```
ggscatter(BMR_b, x = "logBMR", y = "chicks", 
          add = "reg.line", conf.int = TRUE, 
          cor.coef = TRUE, cor.method = "pearson",
          xlab = "(log)BMR", ylab = "Number of chicks")
```

```
ggscatter(BMR_b, x = "logBMR", y = "date", 
          add = "reg.line", conf.int = TRUE, 
          cor.coef = TRUE, cor.method = "pearson",
          xlab = "(log)BMR", ylab = "laying date")
```

```
ggscatter(BMR_b, x = "resBMR", y = "laying date", 
          add = "reg.line", conf.int = TRUE, 
          cor.coef = TRUE, cor.method = "pearson",
          xlab = "mass-independent BMR", ylab = "laying date")
```

```
ggscatter(Data_FFM, x = "resBMR_FFM", y = "resMsum_FFM", 
          add = "reg.line", conf.int = TRUE, 
          cor.coef = TRUE, cor.method = "pearson",
          xlab = "mass-independent BMR", ylab = "mass-independent Msum")
```

```
BMR_mod_FFM <- lm(resBMR_FFM ~ season+sex, data=Data_FFM)
summary(BMR_mod_FFM)
```

```
## 
## Call:
## lm(formula = resBMR_FFM ~ season + sex, data = Data_FFM)
## 
## Residuals:
##      Min       1Q   Median       3Q      Max 
## -0.31889 -0.06418  0.02530  0.08954  0.21435 
## 
## Coefficients:
##             Estimate Std. Error t value Pr(>|t|)
## (Intercept)  0.04720    0.06519   0.724    0.483
## seasonw     -0.11731    0.09219  -1.272    0.227
## sexM         0.09302    0.09219   1.009    0.333
## 
## Residual standard error: 0.1458 on 12 degrees of freedom
##   (1 observation deleted due to missingness)
## Multiple R-squared:  0.1307, Adjusted R-squared:  -0.01418 
## F-statistic: 0.9021 on 2 and 12 DF,  p-value: 0.4315
```

```
Msum_mod_FFM <- lm(resMsum_FFM ~ season+sex, data=Data_FFM)
summary(Msum_mod_FFM)
```

```
## 
## Call:
## lm(formula = resMsum_FFM ~ season + sex, data = Data_FFM)
## 
## Residuals:
##      Min       1Q   Median       3Q      Max 
## -0.14395 -0.09209  0.00000  0.01682  0.29410 
## 
## Coefficients:
##             Estimate Std. Error t value Pr(>|t|)
## (Intercept) -0.19277    0.13885  -1.388    0.202
## seasonw      0.17403    0.15210   1.144    0.286
## sexM         0.07604    0.08782   0.866    0.412
## 
## Residual standard error: 0.1388 on 8 degrees of freedom
##   (5 observations deleted due to missingness)
## Multiple R-squared:  0.264,  Adjusted R-squared:  0.08004 
## F-statistic: 1.435 on 2 and 8 DF,  p-value: 0.2934
```

```
residuals_BMR_mod_FFM <- resid(BMR_mod_FFM)
hist(residuals_BMR_mod_FFM)
```

```
qqnorm(residuals_BMR_mod_FFM)
qqline(residuals_BMR_mod_FFM)
```

```
shapiro.test(residuals_BMR_mod_FFM)
```

```
## 
##  Shapiro-Wilk normality test
## 
## data:  residuals_BMR_mod_FFM
## W = 0.95339, p-value = 0.5794
```
