## Supplementary material for "Seasonal variation in great tit (*Parus major*) energy requirements: reallocation versus increased demand": Supplemetary material

| **Author(s)** | **FMR* (kJ/day)** | **Body mass (g)** | **Mass-specific FMR** | **Period** | **Sample size *n*** |
|---|---|---|---|---|---|
| Sanz, 1998 | 103.20 | 17.5 | 5.90 | Feeding chicks | 10 |
| Tinbergen & Dietz, 1994 | 95.10 | 17.7 | 5.37 | Feeding chicks | 32 |
| Sanz et al., 2000 | 72.00 | 17.4 | 4.14 | Feeding chicks | 7 |
| | 97.90 | 17.7 | 5.53 | | 27 |
| | 103.20 | 17.8 | 5.80 | | 10 |
| Tinbergen & Verhulst, 2000 | 88.70 | 17.4 | 5.10 | Feeding chicks | 13 |
| | 102.10 | 18.0 | 5.67 | | 14 |
| | 105.50 | 17.5 | 6.03 | | 11 |
| De Heij et al., 2008 | 79.30 | 20.1 | 3.95 | Incubation | 14 |
| Bryan & Bryant, 1999 | 111.20 | 21.5 | 5.17 | Incubation | 8 |
| Tinbergen & Wiersma, 2003 | 77.50 | 17.5 | 4.43 | Feeding chicks | 10 |
| | 84.20 | 18.0 | 4.68 | | 10 |
| Verhulst & Tinbergen, 1997 | 92.40 | 17.8 | 5.19 | Feeding chicks | 4 |
| | 90.00 | 17.8 | 5.06 | | 5 |
| | 82.40 | 17.8 | 4.63 | | 9 |
| | 72.60 | 17.7 | 4.10 | | 6 |
| | 92.70 | 17.6 | 5.27 | | 6 |
| | 85.80 | 17.8 | 4.82 | | 5 |
| | 105.60 | 17.5 | 6.03 | | 4 |
| | 86.30 | 17.4 | 4.96 | | 5 |
| | 97.50 | 17.4 | 5.60 | | 7 |
| | 103.10 | 17.6 | 5.86 | | 5 |
| | 88.60 | 17.2 | 5.15 | | 4 |
| | 100.20 | 17.5 | 5.73 | | 6 |
| Nagy et al., *1999* | 96.20 | 18.1 | 5.31 | Winter | 13 |
| *Weighted means* | *93.28* | *17.9* | *5.18* | | |
| Observed in this study | 66.32 | 16.8 | 3.96 | Winter | 2 |
| | 72.71 | 16.3 | 4.48 | Feeding chicks | 3 |
| *Weighted means* | *70.15* | *16.5* | *4.27* | | |

Table S1. Field metabolic rates (FMR), body mass, and mass-specific FMR (FMR divided by the body mass) of studies on great tits (*Parus major*). Weighted mean averages for comparison with the FMR values of this study.

*The use of different $CO_2$ coefficients can significantly affect FMR values, making comparisons a bit difficult.

| Author(s) | BMR (ml $O_2$/min) | Body mass (g) | Mass-specific BMR | Period | Sample size n |
|---|---|---|---|---|---|
| Broggi et al., 2019 | 1.1 | 19.7 | 0.056 | Winter | 159 |
| Playà-Montmany et al., 2021 | 0.96 | 16.6 | 0.058 | - | 24 |
| Ots et al., 2001 | 1.25 | 18.7 | 0.067 | Winter | 42 |
| Daan et al., 1990 | 0.9 | 16 | 0.056 | - | 22 |
| Broggi et al., 2007 | 1.15 | 18.9 | 0.061 | Winter | 324 |
| Broggi et al., 2022 | 1.28 | 19.2 | 0.067 | Winter | 160 |
| Mathot et al., 2016 | 0.96 | 18 | 0.053 | Winter | 111 |
| Reinertsen & Haftorn, 1986 | 1.67 | 16.8 | 0.099 | Winter | 3 |
| Broggi et al., 2004 | 1.15 | 18.5 | 0.062 | Winter | 17 |
| | 1 | 18.8 | 0.053 | | 24 |
| Hissa & Palokangas, 1970 | 1.14 | 18.4 | 0.062 | Winter | 5 |
| | 1.12 | 19 | 0.059 | Summer | 5 |
| Lindström & Kvist, 1995 | 0.77 | 14.9 | 0.052 | Autumn | 1 |
| Steen, 1958 | 1.21 | 18.6 | 0.065 | Winter | 2 |
| Gavrilov, 2014 | 0.98 | 16.4 | 0.06 | Summer | 20 |
| | 1.11 | 17.1 | 0.065 | Winter | 20 |
| Bouwhuis et al., 2011 | 1.36 | 18.2 | 0.075 | Winter | 694 |
| Mathot et al., 2015 | 1.11 | 18.1 | 0.061 | Winter | 142 |
| Broggi & Nilsson, 2023 | 0.89 | 17.6 | 0.051 | Winter | 42 |
| Nilsson & Råberg, 2001 | 0.93 | 15.9 | 0.059 | Winter | 11 |
| | 1.05 | 16.5 | 0.064 | Nest building | 9 |
| | 1.19 | 18.2 | 0.065 | Egg laying | 12 |
| | 1.12 | 16.1 | 0.07 | Feeding chicks | 14 |
| *Weighted means* | *1.21* | *18.4* | *0.066* | | |
| Observed in this study | 1.17 | 16.4 | 0.071 | Winter | 40 |
| | 1.35 | 15.9 | 0.085 | Feeding chicks | 10 |
| *Weighted means* | *1.2* | *16.3* | *0.074* | | |

Table S2. Basal metabolic rates (BMR), body mass, and mass-specific BMR (BMR divided by the body mass) of studies on great tits (*Parus major*). Weighted mean averages for comparison with the BMR values of this study.
